# Cross-membrane cooperation among bacteria can facilitate intracellular pathogenesis

**DOI:** 10.1101/2025.02.09.637186

**Authors:** Daniel Schator, Naren G Kumar, Samuel Joseph U Chong, Timothy K Jung, Eric Jedel, Benjamin E Smith, David J Evans, Suzanne M J Fleiszig

## Abstract

*Pseudomonas aeruginosa* is a Gram-negative opportunistic pathogen able to cause life- and sight-threating infections. Once considered an extracellular pathogen, numerous studies have shown it can survive intracellularly. Previously, we showed that *P. aeruginosa* inside cells can diversify into distinct subpopulations in vacuoles and the cytoplasm. Here, we report that the transition from vacuoles to cytoplasm requires collaboration with the extracellular subpopulation, through Ca^2+^ influx enabled by their type III secretion system (T3SS) translocon pore proteins. Moreover, we show that collaboration among *P. aeruginosa* subpopulations can contribute to disseminating intracellular bacteria *in vivo* in a mouse infection model. This study provides the basis for future studies to investigate how cooperation of extracellular and intracellular bacteria within the host may contribute to disease progression and persistence.

## Introduction

*Pseudomonas aeruginosa* is an opportunistic bacterial pathogen able to cause a wide range of infections, including complication-prone burn wound infections, life-threatening lung infections, and potentially blinding infections of the cornea (*1–6*). The diversity of these infection sites highlights one of the biggest challenges in treating *P. aeruginosa* infections, the adaptability and resilience of this bacterium to survive and thrive under rapidly changing environmental conditions (*7–12*). While *P. aeruginosa* is often labeled as an extracellular pathogen, many investigators have confirmed that clinical and other isolates of *P. aeruginosa* can invade a range of cell types *in vitro* and *in vivo* and are able to establish a complex intracellular lifestyle (*13–22*).

We and others have shown the importance of the bacterial type-III secretion system (T3SS) in intracellular survival by *P. aeruginosa* (*15*, *23*, *24*) and that mutants lacking the T3SS are found only in vacuoles not in the cytoplasm. This has led to the assumption that vacuolar bacteria use these factors to escape vacuoles (*23–26*). Recently we reported that the intracellular lifestyle of *P. aeruginosa* includes diversification into two subpopulations; a rapidly dividing and motile cytoplasmic population that expresses the T3SS, and a stationary and slowly replicating vacuolar population more tolerant to the cell permeable antibiotic ofloxacin (*16*). How *P. aeruginosa* establishes these separate niches and whether one is a precursor to the other is not known.

Here, we aimed to fill this gap in our understanding of *P. aeruginosa* pathogenesis, testing the hypothesis that there are separate pathways for *P. aeruginosa* invasion leading to vacuoles and the cytoplasm, the latter requiring the T3SS. The methods leveraged novel strategies for imaging and image analysis enabling cellular pathogenesis to be studied quantitatively and comparatively *in vitro* and *in vivo*. The results disproved the hypothesis by showing that *P. aeruginosa* enters vacuoles exclusively during the first hour of infection and depends on the extracellular population to access the cytoplasm. Rescue by the extracellular bacteria was found to require their T3SS, specifically the translocon pore proteins via increasing intracellular calcium, which if induced pharmacologically could also trigger vacuolar escape. Surprisingly, the contribution made by T3SS exotoxins was minor. High-resolution confocal imaging of *in vivo* corneal infections confirmed that cooperation among bacterial subpopulations could promote escape from vacuole-like intracellular structures requiring the presence of T3SS^ON^ bacteria, and that it facilitated *in vivo* intracellular pathogenesis. The cross-membrane co-operation among bacterial subpopulations demonstrated here is a novel concept that broadens our understanding of bacterial pathogenesis *in vitro* and *in vivo*.

## Results

### Shorter invasion times produced only vacuolar *P. aeruginosa*

When we previously reported that *P. aeruginosa* PAO1 could establish two different intracellular populations within the host cell, one cytoplasmic, the other vacuolar (*16*), we used 3 hour infection assays before adding a non-cell permeable antibiotic to selectively kill extracellular bacteria. Here, we monitored the fate of bacteria-containing vacuoles arising in these 3 hour infection assays, imaging them over time after extracellular bacteria were killed with the antibiotic. To quantify bacterial vacuoles, we developed a novel image analysis macro described in detail in the methods section. Briefly, the macro detects the signal of fluorescent proteins expressed by the remaining bacteria after extracellular bacteria are killed and classified it into either vacuolar or non-vacuolar signal based on signal area, circularity, and aspect ratio. This was to ensure that only vacuoles supporting viable bacteria were detected, excluding cytoplasmic spreading bacteria. Reflecting the time taken for GFP expression and replication of intracellular bacteria to reach detectable levels after adding arabinose (to induce GFP), there was a gradual increase in vacuole numbers detected during the first few hours after antibiotic addition (**Figure 1A/B**). This was followed by a gradual decrease in the number of vacuoles detected over the 24 hour observation period. Showing that the previously contained bacteria had transitioned to the cytoplasm, they could be seen disseminating within the cell, and the percentage of cells containing intracellular bacteria did not change (**Figure 1C).** Continued use of the non-cell permeable antibiotic throughout the assay ensured that cells could no longer be infected from an extracellular location.

**Figure 1:**
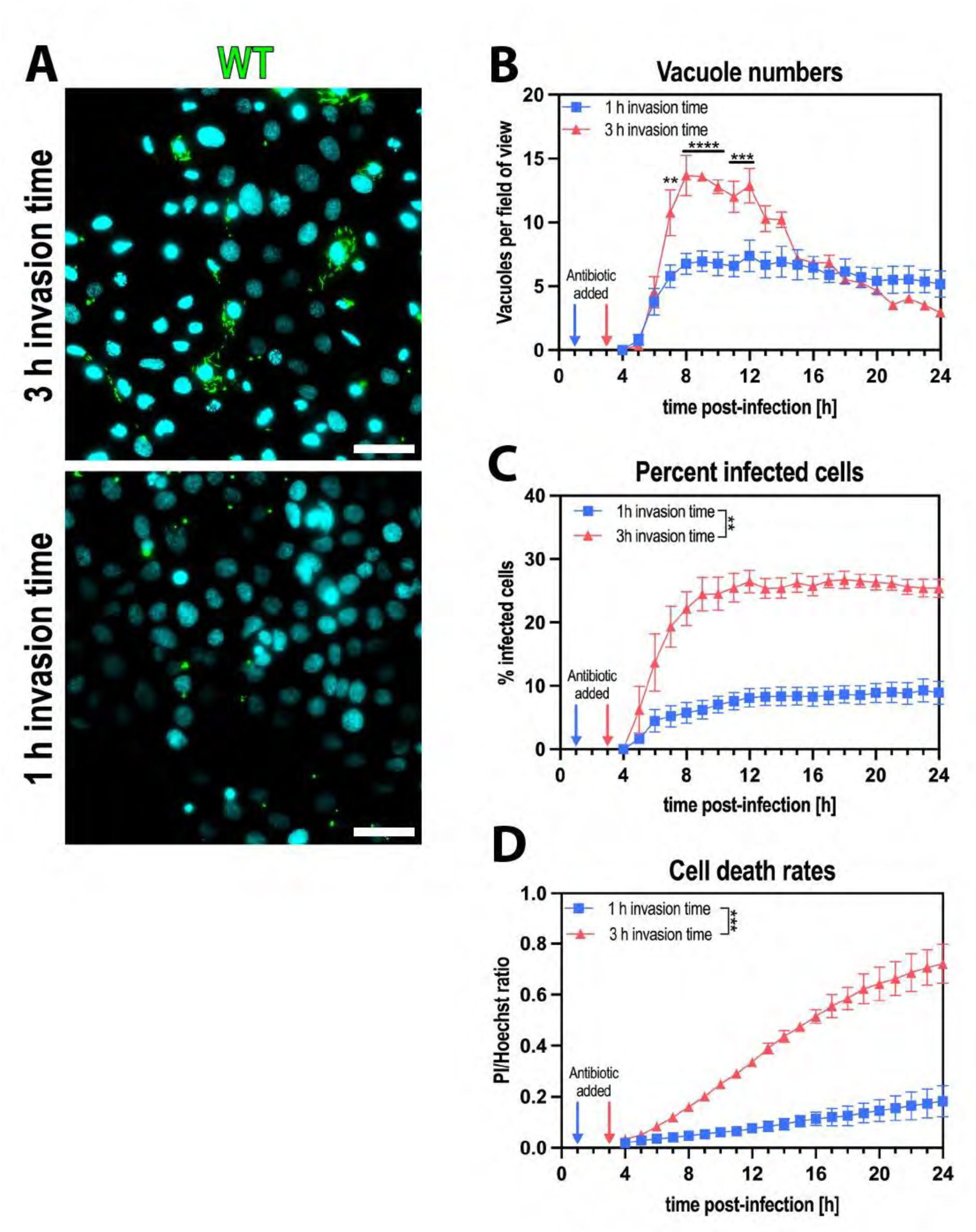
Shorter invasion times produced only vacuolar *P. aeruginosa*. (A) Representative widefield microscopy images of human corneal epithelial cells (hTCEpi) infected with *P. aeruginosa* wildtype at MOI 10 with different invasion times. Images were taken 10 h post-infection at 40x magnification. *P. aeruginosa* (green), Hoechst (cyan). Scale bar equals 50 µm. (B/C/D) Image analysis of timelapse images measuring vacuole numbers (B), percent infected cells (C), and cell death rates (D). Data represented as mean ± SEM, N=3. For statistical analysis, a Two-way ANOVA with multiple comparisons was performed. P ≤ 0.01 = **, P ≤ 0.001 = ***, P ≤ 0.0001 = ****

To understand early events in the sequence leading to the above phenotype, the extracellular bacteria were killed sooner (after 1 hour of infection rather than at 3 hours). Doing so reduced the time allowed for extracellular bacteria to invade cells and the time for them to influence other outcomes. This resulted in fewer infected cells, fewer bacteria-containing vacuoles, and less cell death (**Figure 1B/C/D**). As for 3 hour invasion assays, the number of vacuoles detected steadily increased for ∼ 4 hours after antibiotic addition. However, instead of gradually decreasing over time, vacuole numbers remained stable for the entire 24 hour subsequent observation period, and cytoplasmic bacterial spreading was not observed (**Figure 1A/B**). Thus, bacteria invading during the first hour enter vacuoles only, and are unable to leave if extracellular bacteria are then eliminated.

### Vacuolar release was independent of bacterial numbers

Next, we asked why the fate of bacteria-containing vacuoles inside a cell depended on the length of time that extracellular bacteria were present? We first hypothesized that it was due to replication increasing the inoculum in the longer assays. Indeed, growth curve assays showed efficient bacterial division in the cell culture media between 1 and 3 hours, resulting in a ∼ 5-fold increase in OD_600_ (**Figure 2A**). For some other pathogens, multiplicity of infection (MOI) is thought to impact intravacuolar behavior (*27*). When we tested the impact of increasing the MOI in the shorter 1 hour invasion assays, vacuole numbers, percent infected cells, and cell death all increased in an inoculum-dependent manner (**Figure 2B/C/D**; 2B/D, statistical analysis represented as colored blocks within the graph). Nevertheless, the peak in vacuole numbers occurred at the same time-point for all MOI (∼ 4 hours) and remained stable for the duration of the assay irrespective of MOI (**Figure 2B**). The lack of cytoplasmic disseminating populations was also independent of MOI (**Figure 2E**). Thus, the 5-fold replication occurring during the additional 2 hours of incubation with extracellular bacteria does not on its own explain why vacuole escape occurs after a 3 hour invasion assay and not after a 1 hour invasion assay.

**Figure 2:**
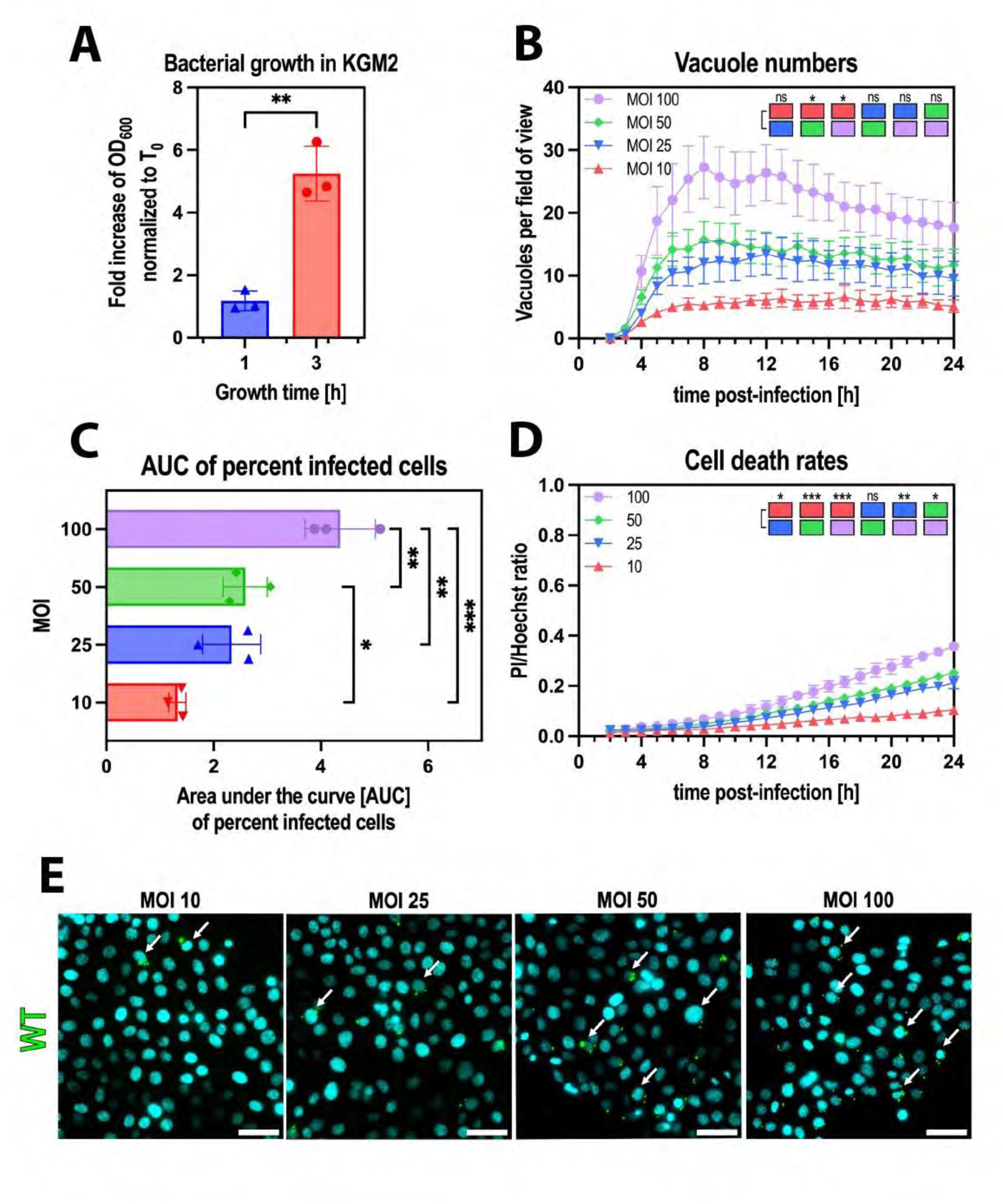
Vacuolar release was independent of bacterial numbers. (A) Quantification of OD_600_ changes of *P. aeruginosa* PAO1 grown in KGM2 for 3 hours at 37°C, without agitation. Values are normalized to T0. (B/C/D) Image analysis of timelapse images measuring vacuole numbers (B), percent infected cells, represented as Area under the curve (AUC) bar plots (C), and cell death rates (D). (E) Representative widefield microscopy images of human corneal cells (hTCEpi) infected with *P. aeruginosa* wild type at different MOIs (10, 20, 50, 100) with 1 h invasion time. Images were taken 10 h post-infection at 40x magnification. *P. aeruginosa* (green), Hoechst (cyan). Arrows highlighting some cells with vacuolar bacteria. Scale bar equals 50 µm. Data represented as mean ± SD (A/C) or SEM (B/D), N=3. For statistical analysis, a Student’s t-test (A), a Two-way ANOVA with multiple comparisons (B/D), or a One-way ANOVA (C) was performed. P ≤ 0.05 = *, P ≤ 0.01 = **, P ≤ 0.001 = ***

### Constitutive expression of the T3SS enabled vacuolar escape in 1 hour invasion assays

We next tested the hypothesis that changes to bacterial gene expression were occurring between 1 hour and 3 hours of an invasion assay which altered the fate of invading *P. aeruginosa*. We focused on the type-III secretion system (T3SS) because it is induced by host cell contact (*28*), known to impact *P. aeruginosa* invasion and intracellular survival (*15*, *22*, *23*, *29*), and because cells infected with mutants lacking various components of the T3SS do not escape vacuoles even in 3 hour invasion assays(*13*). Thus, we compared mutants in specific T3SS genes that toggle T3SS expression off and on, including a Δ*exsA* (T3SS^OFF^) mutant lacking the transcriptional activator of all T3SS genes, and a Δ*exsE* (T3SS^ON^) mutant constitutively expressing the T3SS because ExsE otherwise represses the activator ExsA (*30*). Both mutants differ from wild-type *P. aeruginosa* which requires induction by host cell contact or low calcium conditions to express the T3SS, which for wild-type occurs in only a fraction of the population due to the bistability of T3SS gene expression. Constitutive expression of the T3SS by the Δ*exsE* mutant (T3SS^ON^), and not by wild-type or the Δ*exsA* mutant (T3SS^OFF^), was confirmed in both T3SS-inducing and non-inducing conditions using a T3SS reporter plasmid (pJNE05) (**Figure 3A**).

**Figure 3:**
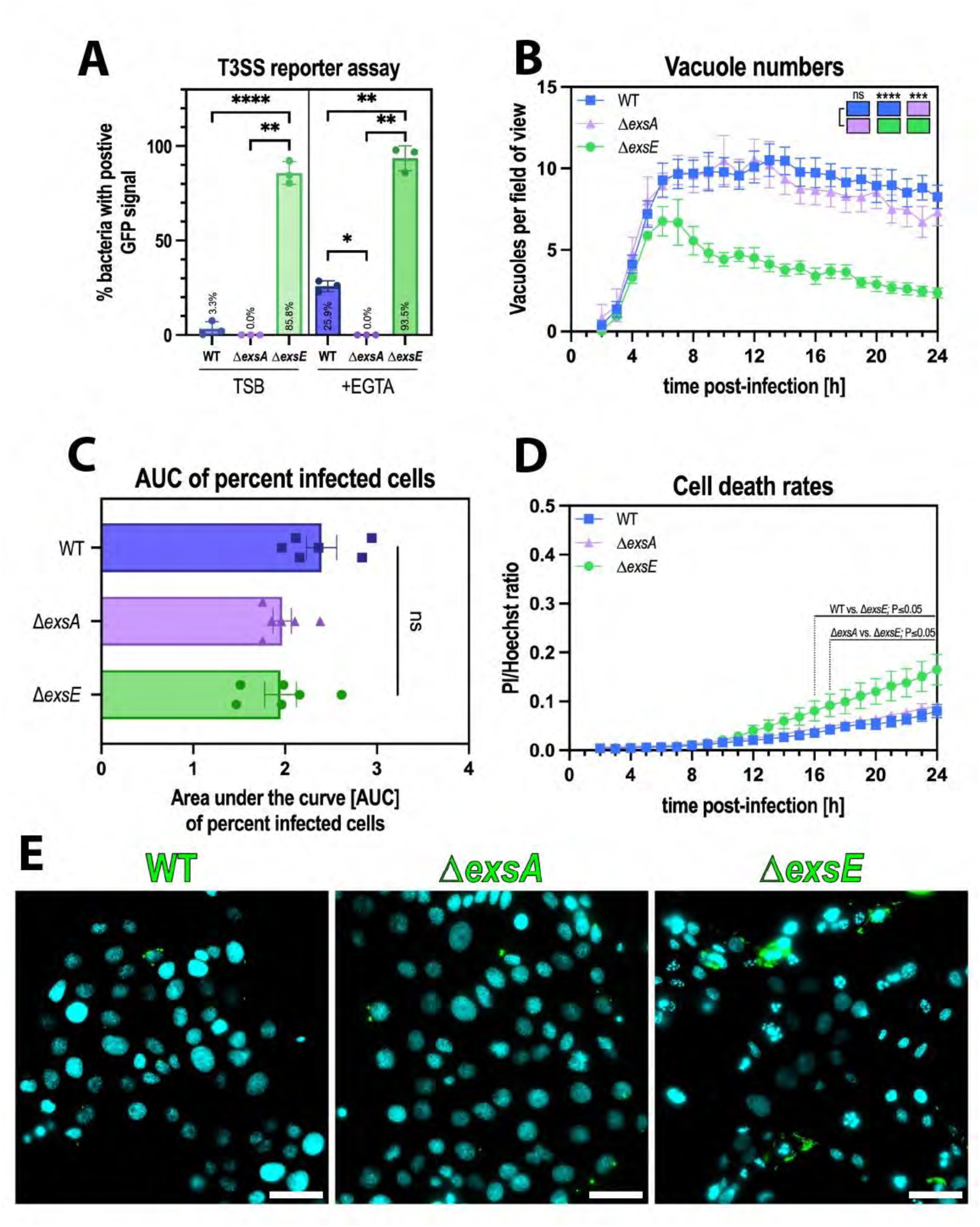
Constitutive expression of the T3SS enabled vacuolar escape in 1 hour invasion assays. (A) Quantification of T3SS-expressing (GFP positive) bacteria in a spotting assay comparing WT, Δ*exsA*, and Δ*exsE* under non-inducing (TSB) and inducing (+EGTA) conditions. (B/C/D) Image analysis of timelapse images measuring vacuole numbers (B), percent infected cells, represented as Area Under the Curve (AUC) bar plots (C), and cell death rates (D). (E) Representative widefield microscopy images of human corneal epithelial cells (hTCEpi) infected with *P. aeruginosa* wildtype, Δ*exsA* (T3SS^OFF^) or Δ*exsE* (T3SS^ON^) strains at MOI 50 using 1 h invasion time. Images were taken at 10 hours postinfection at 40x magnification. *P. aeruginosa* (green), Hoechst (cyan). Scale bar equals 50 µm. Data represented as mean ± SD (A/C) or SEM (B/D), N=3 (A), N=8 (B/D), N=6 (C). For statistical analysis, a One-way ANOVA with multiple comparisons (A/C) or Two-way ANOVA with multiple comparisons (B/D) was performed. P ≤ 0.01 = **, P ≤ 0.001 = ***, P ≤ 0.0001 = ****

Results showed that T3SS^ON^ and T3SS^OFF^ mutants invaded a similar percentage of cells as wild-type in the1 hour infection assays and yielded a similar number of vacuoles (**Figure 3B/C**; 3B, statistical analysis represented as colored blocks). Following that, the results differed for the T3SS^ON^ mutants. Rather than remaining stable, there was a decline in vacuole numbers corresponding with bacterial dissemination through the cell cytoplasm, and higher levels of cell death several hours after vacuolar release was observed (**Figure 3B/D/E**). This suggests that T3SS expression status could determine vacuolar release in the short (1 hour) invasion assays.

### Extracellular bacteria enabled release of vacuolar populations in a T3SS-dependent manner

Having observed that T3SS expression status is important for vacuolar release and that wild-type vacuole-contained bacteria were unable to trigger their own release without assistance from extracellular bacteria after 1 hour, we tested the hypothesis the extracellular bacteria use their T3SS to rescue them. The rationale was that wild-type bacteria needed longer invasion times to trigger vacuolar release, whereas the T3SS^ON^ mutant did not, potentially reflecting time needed for the T3SS to be induced in extracellular bacteria upon host cell contact or for injection of responsible effectors.

To test this, we asked if co-infection with the T3SS^ON^ mutants (constitutively expressing the T3SS) could rescue wild-type and/or T3SS^OFF^ mutants from vacuoles they were otherwise trapped in. To distinguish mutants/wild-type from one another, we used enhanced Green Fluorescent Protein (eGFP) (green) and mScarlet3 (red) expression under the control of an arabinose-inducible promoter.

The results showed that co-infection with T3SS^ON^ bacteria could rescue both wild-type and T3SS^OFF^ bacteria from their vacuoles in the 1 hour invasion assays. This was illustrated by a decrease in the number of vacuoles over time during continued incubation in antibiotic (**Figure 4A/B**, statistical analysis represented as colored blocks), along with the appearance, replication and dissemination of bacteria in the cell cytoplasm compared to when T3SS^ON^ bacteria were not present (**Figure 4C**). This occurred without impact on the percentage of cells containing intracellular wild-type or T3SS^OFF^ bacteria, showing that the mechanism did not involve loss of cells or cell viability (**Figure 4D/E**). While cell death was impacted by the presence of the Δ*exsE* strain, this occurred several hours after the drop in vacuole numbers (**Figure 4F/G**).

**Figure 4:**
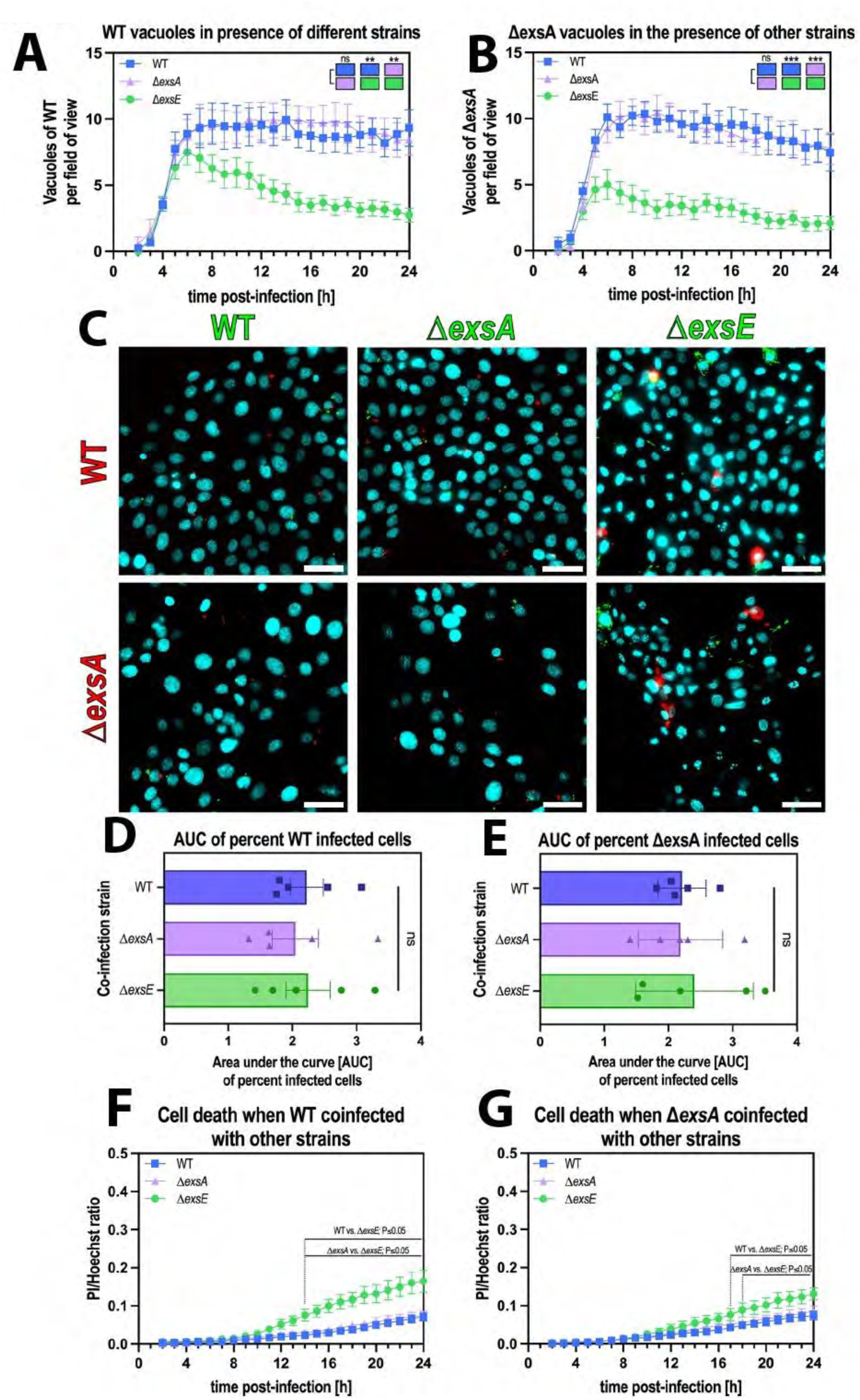
Extracellular bacteria enable release of vacuolar populations in a T3SS-dependent manner. (A/B) Image analysis of timelapse images measuring vacuole numbers for WT (A) or Δ*exsA* (B) when co-infected with different strains. (C) Representative widefield microscopy images of human corneal epithelial cells (hTCEpi) infected with *P. aeruginosa* wild-type/Δ*exsA* (red) co-infected with different strains (wild-type, Δ*exsA*, Δ*exsE*; all in green) at MOI 100 (MOI 50 per strain). Images were taken 10 h post-infection at 40x magnification. *P. aeruginosa* (green or red), Hoechst (cyan). Scale bar equals 50 µm. (D/E) Area under the curve bar plots for the percent infected cells of red WT (D) or Δ*exsA* (E) bacteria in presence of other strains. (F/G) Image analysis of timelapse images measuring cell death rates in WT (F) or Δ*exsA* (G) infections, coinfected with other strains. Data represented as mean ± SEM (A/B/F/G) or SD (D/E), N=8 (A/B), N=5 (C/D), N=6 (F/G). For statistical analysis, a Two-way ANOVA with multiple comparisons (A/B/F/G) or a One-way ANOVA with multiple comparisons (D/E) was performed. P ≤ 0.01 = **, P ≤ 0.001 = ***

Having shown that cooperation between T3SS^ON^ and T3SS^OFF^ populations can allow *P. aeruginosa* to escape to the cytoplasm, we next asked if the extracellular population was responsible for the rescue phenotype, the alternative being that the responsible T3SS^ON^ bacteria had invaded the same cell. To explore this directly, we quantified the incidence of double invasion, i.e. T3SS^ON^ and T3SS^OFF^ or wild-type being inside the same cell, comparing predicted values based on probability calculations with observed values, which showed no significant differences (**Table 1**). We then compared the predicted values of vacuolar release (i.e. if double invasion of the same cell is necessary for rescue) with the observed values.

The calculations revealed an almost 10-fold higher incidence of vacuolar release for both wild-type and Δ*exsA* (T3SS^OFF^) mutants than was predicted if double infection of the same cell was sufficient to enable release (**Table 1**). This outcome supported the concept that invasion of the same cell was not required for T3SS-expressing *P. aeruginosa* to release their colleagues from vacuoles. Since Δ*exsE* (T3SS^ON^) mutants rescued Δ*exsA* (T3SS^OFF^) mutants, it follows that it was the T3SS of the “rescuing” not “rescued” bacteria that contributed. Together, these results implicate the T3SS of the extracellular population in triggering release of intracellular *P. aeruginosa* from their vacuoles, thereby enabling them to escape into the cell cytoplasm.

**Table 1:** Probability of double invasion and vacuolar release.

| Strains | Infection rate of single strains (Mean ± SD) | Double infection rate with ΔexsE (Mean ± SD) | | | Rate of vacuolar release in WT/ΔexsA infected cells (Mean ± SD) | | |
| --- | --- | --- | --- | --- | --- | --- | --- |
|  |  | Predicted | Observed | Significance | Predicted (if double invasion is necessary for release) | Observed | Significance |
| WT | 12.11 % (±4.73) | 1.08 % (±0.72) | 0.56% (±0.25) | ns | 4.70 % (±1.65) | 45.30 % (±3.91) | p < 0.0001 |
| ΔexsA | 13.68% (±5.69) | 1.23 % (±0.85) | 0.54% (±0.16) | ns | 4.23 % (±1.34) | 48.23 % (±3.23) | p < 0.0001 |
| ΔexsE | 8.01% (±3.12) | ND | ND | ND | ND | ND | ND |

### Triggering of vacuolar escape depended on T3SS translocon pore proteins

We and others have shown that T3SS components can contribute to the intracellular lifestyle of *P. aeruginosa*, most importantly ExoS and the translocon pore proteins. To determine which T3SS component(s) participate in rescue from vacuoles, we constructed Δ*exsE* mutants in strains with mutations in the T3SS machinery (i.e. needle and translocon) as well as exotoxins, to explore contributions of T3SS components under conditions that ensured consistent expression of the T3SS machinery. This included Δ*exsE*Δ*pscC*, Δ*exsE*Δ*popBD* and Δ*exsE*Δ*exoS*/*T*/*Y* mutants (lacking the T3SS needle, translocon pore proteins and known exotoxins, respectively), which were then used for co-infection experiments with wild-type and T3SS^OFF^ (Δ*exsA*) mutants.

The results showed that in the absence of the needle (Δ*pscC*) or the translocon pore (Δ*popBD*), T3SS^ON^ mutants lost their ability to rescue vacuolar wild-type or T3SS^OFF^ (Δ*exsA*) mutants through co-infection (**Figure 5A/B**, statistical analysis represented as colored blocks). This did not impact percent invaded cells in any instance (**Figure 5C/D**).

**Figure 5:**
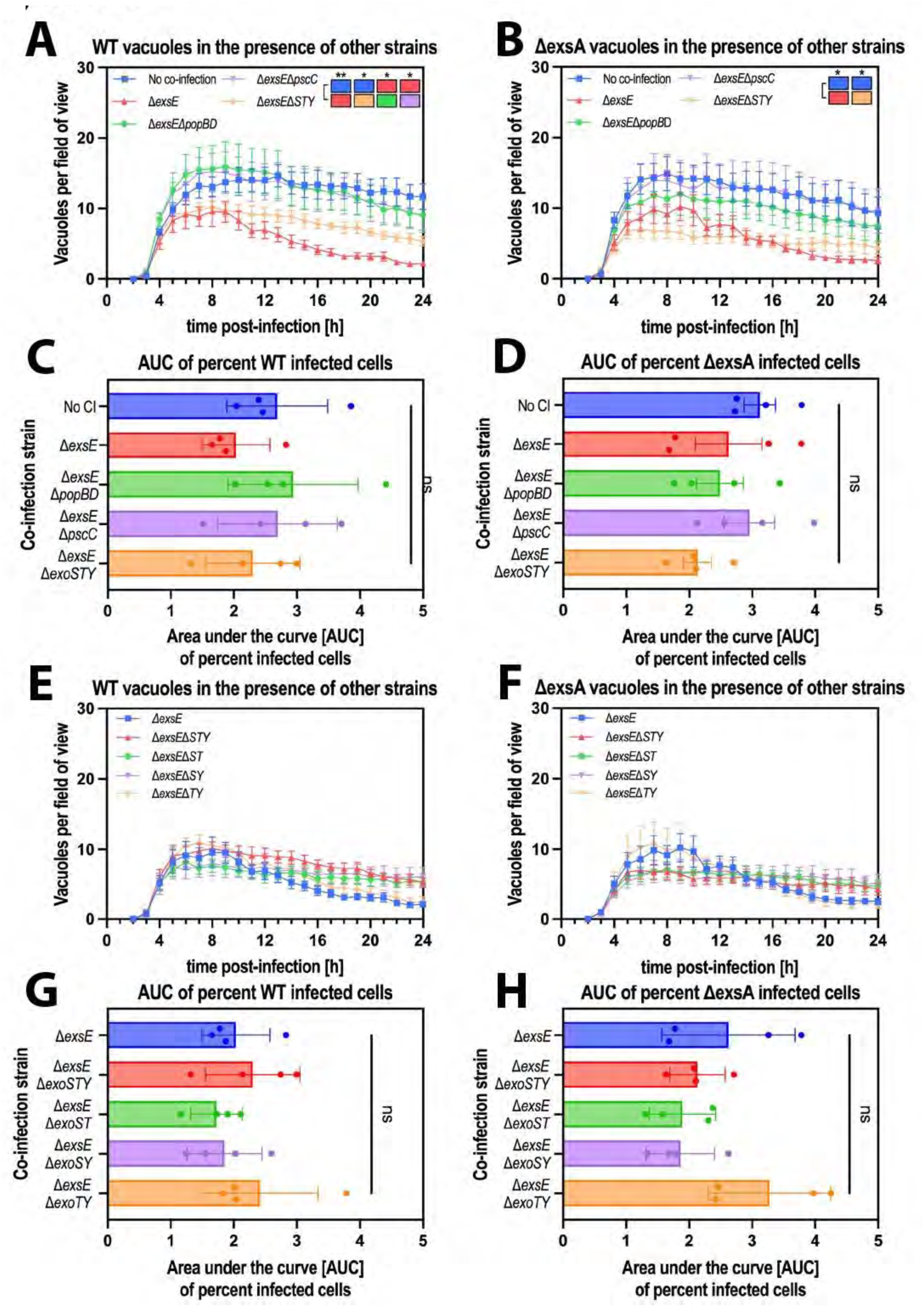
Triggering of vacuolar escape depended on T3SS translocon pore proteins. (A/B/E/F) Image analysis of timelapse images measuring vacuole numbers for WT (A/E) or Δ*exsA* (B/F) in human corneal epithelial cells (hTCEpi) at MOI 50 when co-infected with different strains. (C/D/G/H) Area Under the Curve bar plots for the percent cells infected with WT (C/G) or Δ*exsA* (D/H) in presence of other strains. Data represented as mean ± SEM (A/B/E/F) or SD (C/D/G/H), N=4. For statistical analysis, a Two-way ANOVA with multiple comparisons (A/B/E/F) or a One-way ANOVA with multiple comparisons (C/D/G/H) was performed. P ≤ 0.05 = *, P ≤ 0.01 = **

Surprisingly, the mutant lacking T3SS exotoxins encoded by strain PAO1 (Δ*exoS*/*T*/*Y*) caused only a partial loss of the phenotype that was not statistically significant (**Figure 5A/B**). To account for the possibility of exotoxins having opposing impacts on the phenotype, we tested them individually using double exotoxin knockouts - again in the background of the Δ*exsE* to enable constitutive expression of the T3SS (Δ*exsE*Δ*exoS*/*T*, Δ*exsE*Δ*exoS*/*Y,* Δ*exsE*Δ*exoT*/*Y*). Only the Δ*exsE*Δ*exoT*/*Y* mutant (i.e. still able to express *exoS*) showed a phenotype similar to the Δ*exsE* knockout, however the difference to the other double knockouts was not statistically significant (**Figure 5E/F**). Similarly, there was also no impact on percent infected cells (**Figure 5G/H**). Thus, triggering of vacuolar release by the T3SS of extracellular bacteria requires the translocon, with ExoS playing a negligible role.

### Influx of Ca^2+^ promoted release of vacuolar bacteria

The mechanism by which the T3SS translocon proteins of *P. aeruginosa* act on host cells is generally thought to involve pore formation in the host cell membrane, which enables influx of the T3SS exotoxins that are secreted via the T3SS. Recently, it was shown that the PopBD translocon pore can function directly as a pore forming toxin, promoting K^+^-efflux to alter the host epigenome (*31*). Similar effects on intracellular ion levels have been shown for other pore-forming toxins, including listeriolysin O, adenylate cyclase toxin (CyaA), or suilysin (*32–34*). Here, we explored if pore-formation and/or intracellular ion concentration fluxes can rescue *P. aeruginosa* from vacuoles, treating wild-type infected cells with the general pore-forming agent saponin and various ionophores that promote ion transport through cell membranes. The latter included Valinomycin, a K^+^-specific ionophore and calcimycin, an ionophore with high affinity for Ca^2+^. Of these, only calcimycin promoted release of intracellular bacteria from vacuoles, suggesting intracellular Ca^2+^ levels impact vacuole stability (**Figure 6A**, statistical analysis represented as colored blocks). We also assessed if any of the treatments impacted percent infected cells or bacterial growth that might have explained changes in vacuole numbers, which was not the case (**Figure 6B/C**).

**Figure 6:**
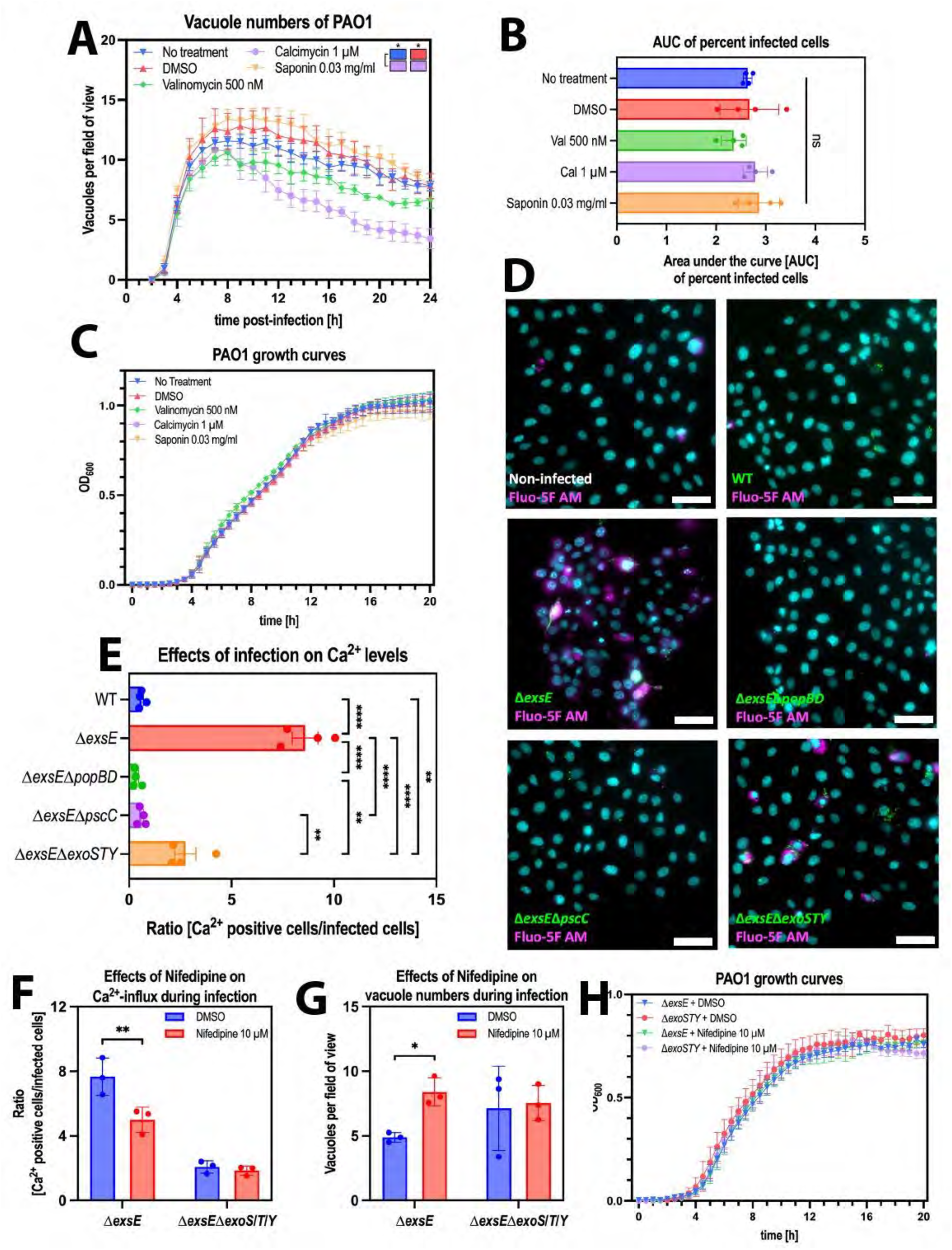
Influx of Ca2+ promoted release of vacuolar bacteria. (A) Image analysis of timelapse images measuring vacuole numbers for WT in corneal epithelial cells (hTCEpi) at MOI 100 when treated with different compounds. (B) Area Under the Curve bar plots for the percent infected cells of wild-type PAO1 in human corneal epithelial cells (hTCEpi) at MOI 100 when treated with different compounds. (C) Growth of PAO1 wild-type measured at OD_600_ in the presence of different compounds. (D) Representative widefield microscopy images of human corneal epithelial cells (hTCEpi) infected with different T3SS mutant strains of PAO1 (green) at MOI 100 and increases in Ca^2+^ levels are detected using Fluo-5F AM (magenta). Images were taken 5 h post-infection at 40x magnification. *P. aeruginosa* (green), Hoechst (cyan), Fluo-5F AM (magenta). Scale bar equals 50 µm. (E) Bar plot of ratio of hTCEpi cells with positive Ca^2+^ signal (Fluo-5F AM) over number of infected cells of the respective strain 5 hours post infection. (F) Ratio of hTCEpi cells with positive Ca^2+^ signal (Fluo-5F AM) 5 hours post-infection over number of infected cells of the respective strain when treated with 10 µM Nifedipine for 20 h prior to infection, compared to DMSO (vehicle control). (G) Number of vacuoles per field of view of the respective strains 5 hours post infection in hTCEpi cells when treated with 10 µM Nifedipine for 20 h prior to infection, compared to DMSO (vehicle control). (H) Growth of PAO1 Δ*exsE* and Δ*exsE*Δ*exoS*/*T*/*Y* measured at OD_600_ in the presence of 10 µM Nifedipine or vehicle control (DMSO). Data represented as mean ± SEM (A) or SD (B/C/E/F/G/H), N=4 (A/B/E) N=3 (C/F/G/H). For statistical analysis, a Two-way ANOVA with multiple comparisons (A/C/F/G/H) or One-way ANOVA with multiple comparisons (B/E) was performed. P ≤ 0.05 = *, P ≤ 0.01 = **, P ≤ 0.001 = ***, P ≤ 0.0001 = ****

We then used a calcium sensor to monitor intracellular calcium levels four hours post-infection using the 1 hour invasion time model comparing T3SS^ON^ (Δ*exsE*) mutants to T3SS^ON^ needle mutants (Δ*exsE*Δ*pscC*) and T3SS^ON^ translocon mutants (Δ*exsE*Δ*popBD*). This early time-point was chosen to assess the Ca^2+^ influx within the first few hours of infection. A T3SS^ON^ exotoxin effector mutant (Δ*exsE*Δ*exoS*/*T*/*Y*) was also included to study the impact of the T3SS machinery (which includes the needle and translocon proteins) in the absence of the T3SS exotoxins. Results showed that intracellular Ca^2+^ levels changed during infection with the Δ*exsE* mutant (constitutive T3SS expression), but not with wild-type *P. aeruginosa* (bistable T3SS expression), Δ*exsE*Δ*popBD* (constitutive T3SS, translocon mutant) or Δ*exsE*Δ*pscC* (constitutive T3SS, needle mutant) infection (**Figure 6D**). The Δ*exsE*Δ*exoS*/*T*/*Y* (T3SS^ON^, no exotoxins) mutant showed an intermediary phenotype, indicating that the T3SS exotoxins play a minor role in Ca^2+^ influx, similar to that noted for vacuolar release in co-infection experiments. Thus, the ability to raise intracellular Ca^2+^ levels mirrored the “vacuole rescue” capacity of the various mutants, both implicating the translocon as a key contributor. This suggests that host cell invasion was not required to raise intracellular Ca^2+^ levels, the number of Ca^2+^-positive cells far exceeded the number of cells containing intracellular bacteria; being 8-fold higher for T3SS^ON^ (Δ*exsE*) mutants and remaining 3-fold higher when they additionally lacked the effectors (Δ*exsE*Δ*exoS*/*T*/*Y*) (**Figure 6E**). Together, these results implicated the T3SS translocon pore proteins of the extracellular population as being largely responsible for triggering Ca^2+^ influx, correlating with their capacity to rescue bacteria from vacuoles inside the cell.

To determine if Ca^2+^ influx driven by the T3SS translocon was dependent on cellular Ca^2+^-channels, cells were treated with the Ca^2+^-channel inhibitor nifedipine (10 µM) before infecting the cells with either the Δ*exsE* (T3SS^ON^) or Δ*exsE*Δ*exoS*/*T*/*Y* (T3SS^ON^, no exotoxin) mutants. The rationale for inclusion of the exotoxin mutant was to again study the phenotype caused by the needle/translocon without impacts of the exotoxins, whereas needle/translocon mutants were <u>not</u> included because they do not promote Ca^2+^ influx. Ca^2+^ influx caused by the T3SS^ON^ (Δ*exsE*) mutant was inhibited by pretreatment with the cellular Ca2+ channel inhibitor nifedipine. In contrast, it had no impact on Ca^2+^ influx caused by the T3SS^ON^ exotoxin (Δ*exsE*Δ*exoS*/*T*/*Y*) mutant that lacks exotoxins while able to express translocon and needle proteins (**Figure 6F**). Together this shows that while the T3SS translocon pore driven Ca^2+^ influx functions independently of cellular Ca^2+^-channels, since it is not inhibited by nifedipine, exotoxin driven Ca^2+^ influx depends on host Ca^2+^-channels and can be inhibited with nifedipine.

After confirming that nifedipine had no impact on bacterial growth (**Figure 6H**), we used it to explore cause and effect relationships between Ca^2+^ influx and vacuole release enabled by T3SS exotoxins. Results showed that blocking cellular Ca^2+^channels with nifedipine could preserve vacuole numbers compared to infection without the drug for T3SS^ON^ (Δ*exsE)* mutants, but not when the bacteria also lacked the T3SS exotoxins (Δ*exsE*Δ*exoS*/*T*/*Y)* (**Figure 6G**). Thus, increasing intracellular Ca^2+^ levels through cellular Ca^2+^ channels can promote vacuolar release driven by T3SS exotoxins. Whether Ca^2+^ influx mediated by the T3SS translocon contributes to how it promotes vacuole release remains to be determined and will require delineating the alternative Ca^2+^ influx pathway and then developing a strategy to block it.

### Bacterial cooperation occurred *in vivo*

The above mechanistic experiments were done using telomerase-immortalized human corneal epithelial cells grown *in vitro*. While these are more likely to be relevant to actual infections than the use of transformed cell lines, factors found *in vivo* can modify host microbe interactions, as we have shown for tear fluid impacts on the corneal epithelium (*35–37*). Thus, we next explored if cooperation between diverse *P. aeruginosa* populations also occurs during *in vivo* infection. For this, we used a murine corneal infection model and utilized live confocal microscopy to determine the spatial and subcellular localization of bacteria. The cornea is uniquely suited to high resolution live imaging due to its natural transparency and the superficial location of its multilayered epithelium.

Briefly, corneas were scratched and infected for 16 hours before extracellular bacteria were killed using Amikacin (300 µg/ml) followed by arabinose induction of GFP fluorescence to enable visualization of only intracellular bacteria. The eyes were enucleated and directly imaged without fixation. Use of different combinations of fluorescent and non-fluorescent bacteria allowed us to determine the impact of bacterial cooperation between the wild-type and different T3SS mutants. Combinations used included the following mutants in the following proportions, with GFP expressed only in the bacteria we were tracking (i.e. only in the to be rescued bacteria and not in the rescuers): 90% Δ*exsA*-GFP+10% Δ*exsA* (T3SS^OFF^+T3SS^OFF^), 90% Δ*exsA*-GFP+10% Δ*exsE* (T3SS^OFF^+T3SS^ON^), as well as 10% Δ*exsE*-GFP (T3SS^ON^) as a low inoculation size control to assess the impact of Δ*exsE* alone.

Infection with only Δ*exsA* (T3SS^OFF^) mutants (90% Δ*exsA*-GFP+10% Δ*exsA*) resulted in few foci of intracellular bacteria, showing clustering within corneal epithelial cells into bigger vacuole-like structures. Co-infection of Δ*exsA* (T3SS^OFF^) with Δ*exsE* (T3SS^ON^) mutants (90% Δ*exsA*-GFP+10% Δ*exsE*) resulted in a very different phenotype for the intracellular T3SS^OFF^ bacteria, which were then able to spread further within the tissue with dispersal of the previously seen vacuole-like structures. This was similar to results with 10% Δ*exsE* (T3SS^ON^) mutants used alone. Thus, the T3SS^OFF^ (Δ*exsA*) bacteria behaved more like T3SS^ON^ bacteria when in the presence of Δ*exsE* bacteria (**Figure 7A**, **Video 1**). Differences in tissue spreading were quantified using histograms of GFP-intensity in 100 µm × 100 µm areas. T3SS^OFF^-GFP+T3SS^OFF^ infections led to a single peak of GFP-signal due to their more localized foci containing intracellular bacteria, whereas 90% Δ*exsA* T3SS^OFF^-GFP+10% Δ*exsE* T3SS^ON^ and 10% Δ*exsE* T3SS^ON^-GFP alone led to a more spread out GFP signal, reflecting spread of the intracellular bacteria throughout the tissue (**Figure 7B**). Furthermore, image analysis revealed that 90% Δ*exsA* T3SS^OFF^-GFP+ 10% Δ*exsE* T3SS^ON^ and the low inoculation size control (10% Δ*exsE* T3SS^ON^-GFP) led to a higher percentage of cells containing intracellular bacteria compared to 90% Δ*exsA* T3SS^OFF^-GFP+10% Δ*exsE* T3SS^OFF^, again pointing towards a more significant intracellular infection spread phenotype (**Figure 7C**). The bacterial aggregates also differed quantitively in their average sizes, those infected with only Δ*exsA* T3SS^OFF^ mutants showing aggregates around twice the size (presumably vacuoles) of average aggregate size when Δ*exsE* T3SS^ON^ mutants were used alone or in combination with Δ*exsA* T3SS^OFF^ mutants (**Figure 7D**). Thus, bacterial cooperation among different bacterial populations also occurs *in vivo* to promote dissemination of intracellular bacteria during *P. aeruginosa* infection, both within individual cells and throughout the tissue.

**Figure 7:**
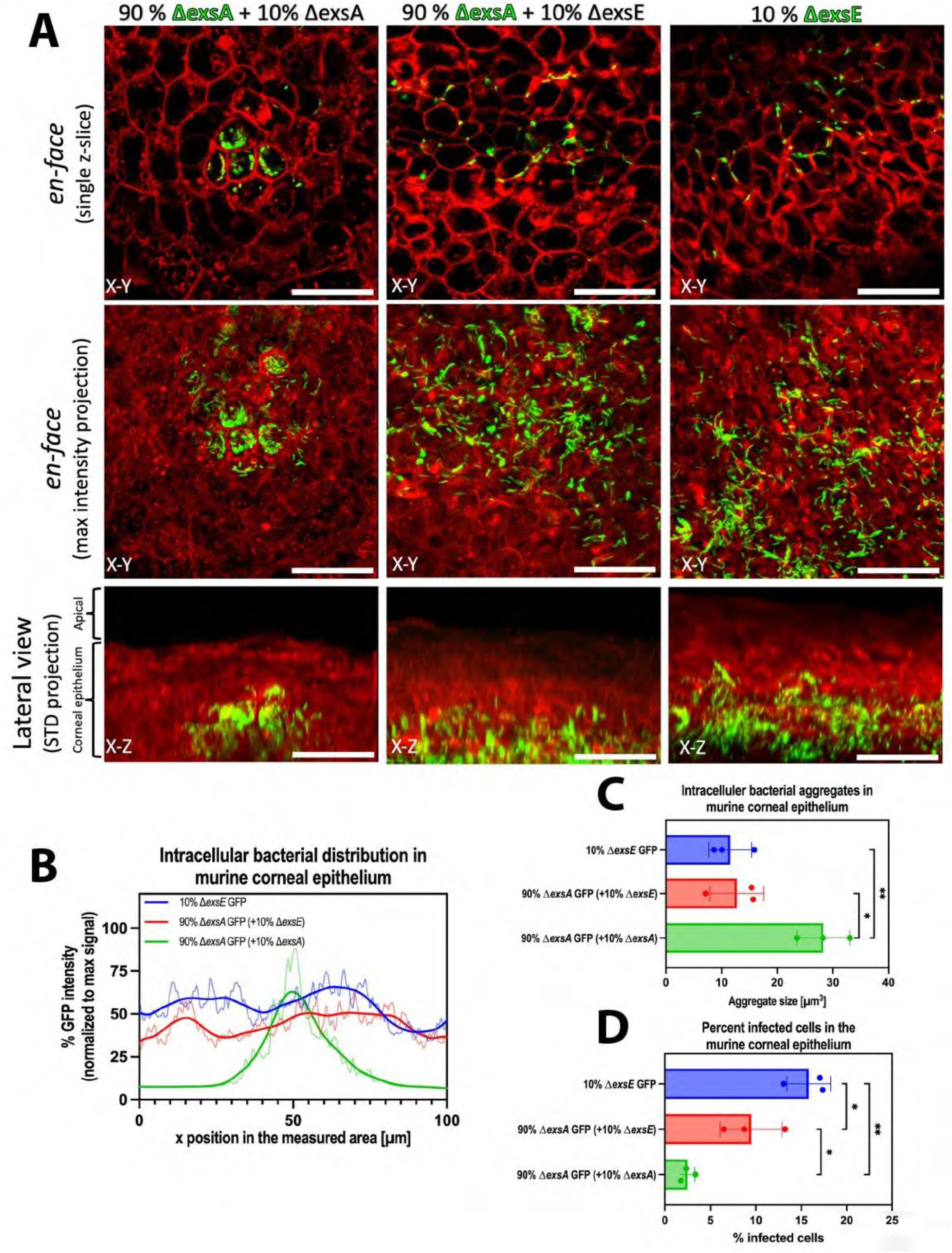
Bacterial cooperation occured *in vivo*. (A) Representative *in vivo* confocal microscopy images of murine corneas (ROSAmT/mG mice, dtTomato cell membranes, red) infected with different combinations of fluorescent (green) and non-fluorescent bacteria (Δ*exsA*-GFP+10% Δ*exsA*, 90% Δ*exsA*-GFP+10% Δ*exsE*, 10% Δ*exsE*-GFP). Images were taken 20 hours post-infection using the murine scratch-infection model, imaging infection foci within the corneal epithelium using a 60x water-immersion objective. Top panel shows *en-face* view of a single z-slice, middle panel shows *en-face* view of max intensity projections, bottom panel shows the corresponding lateral view as STD-projection. Scale bar equals 30 µm (B) Histogram with LOWESS curve of bacterial GFP-signal distribution using different combinations of fluorescent and non-fluorescent bacteria (Δ*exsA*-GFP+10% Δ*exsA*, 90% Δ*exsA*-GFP+10% Δ*exsE*, 10% Δ*exsE*-GFP) normalized to max GFP signal for each image. (C) Bar plots quantifying percent cells infected with GFP-producing bacteria under the three different infection conditions (10% Δ*exsE*, 90% Δ*exsA*-GFP+10% Δ*exsE*, Δ*exsA*-GFP+10% Δ*exsA*). (D) Bar plot quantifying the average size of intracellular GFP-aggregates (particles over 5 μm^3^) in the three infection conditions (10% Δ*exsE*, 90% Δ*exsA*-GFP+10% Δ*exsE*, Δ*exsA*-GFP+10% Δ*exsA*). Data represented as mean ± SD, N=3 (C/D). For statistical analysis, a One-way ANOVA with multiple comparisons (C/D) was performed. P ≤ 0.05 = *, P ≤ 0.01 = **

## Discussion

The goal of this study was to understand the dynamics of intracellular niche formation by *P. aeruginosa*. Since the T3SS is required for *P. aeruginosa* to colonize the cytoplasm of epithelial cells, it was previously assumed that T3SS expression by the intracellular population enables either escape or evasion of vacuoles (*23–25*). Here, we found that during the first hour *P. aeruginosa* enters vacuoles from which they cannot escape if the extracellular population is eliminated using a non-cell permeable antibiotic (**Figure 1A/B**). This does not depend on the MOI or how long they are monitored after antibiotic addition (**Figure 1B**, **Figure 2B/E**). Co-infection experiments using mutants constitutively expressing the T3SS (Δ*exsE*) showed that transition from vacuoles into the cytoplasm required the T3SS to be expressed by extracellular bacteria (**Figure 4A-C**). This was largely attributable to the T3SS translocon with a minor role for the T3SS exotoxins (**Figure 5A/B**). T3SS-dependent induction of Ca^2+^ influx was found to contribute (**Figure 6D/E**), with pharmacologically induced Ca^2+^ influx alone able to trigger vacuolar escape of *P. aeruginosa* (**Figure 6A**). A similar phenotype was found *in vivo* during corneal infection, with the presence of T3SS constitutively expressing bacteria (Δ*exsE*) influencing the location of intracellular T3SS^OFF^ mutants (Δ*exsA*) that otherwise remained in vacuole-like structures in cells and in smaller foci within the infected tissue (**Figure 7A**, **Supp. Video 1**). Cross-membrane co-operation among bacterial subpopulations as demonstrated here is a novel concept that broadens our understanding of bacterial pathogenesis *in vitro* and *in vivo*.

Bistability of T3SS expression creates both T3SS^ON^ and T3SS^OFF^ individuals in any wild-type *P. aeruginosa* population (*23*). Since the T3SS encodes potentially antiphagocytic exotoxins (*23*, *38*) cooperation should allow *P. aeruginosa* significant flexibility as a pathogen. Indeed, the model emerging from our research is that when *P. aeruginosa* encounters a suitable host cell, some members of the population not expressing the T3SS are rapidly engulfed into vacuoles with next steps depending on conditions outside the host cell. If suitable for extracellular bacteria, it could favor transition of the vacuolar population into the cytoplasm, wherein they strongly express the T3SS (*16*) and replicate rapidly (*19*) before being released back into the extracellular space either by egress in plasma membrane blebs (*13*) or when the cell eventually dies at a much later time point (*39*). If conditions are instead unfavorable for extracellular bacteria (e.g. in the presence of antibodies, immune cells, and/or antimicrobial peptides/antibiotics), intracellular bacteria could persist in vacuoles wherein they grow more slowly and are more antibiotic tolerant (*16*), providing a reservoir for infection reseeding at a later time point if extracellular conditions become more suitable.

Finding that extracellular *P. aeruginosa* can contribute to intracellular pathogenesis does not diminish the importance of extracellular pathogenesis as a separate entity. Instead, it reveals an additional role for extracellular bacteria in disease pathogenesis. Extracellular *P. aeruginosa* can use toxins, other secreted factors, and/or surface exposed molecular components/appendages to directly kill mammalian cells, subvert their function, break down tissue barriers, and/or form extracellular biofilms, all of which can contribute to pathogenesis independently of intracellular bacteria (*40–48*). The relative roles of intracellular and extracellular bacteria in driving pathogenesis within the complexity of an *in vivo* system is likely to vary spatially, temporally and conditionally.

The mechanism by which the T3SS of extracellular *P. aeruginosa* promotes release of vacuole-confined colleagues was found to involve Ca^2+^ influx. This was supported by data showing that T3SS translocon-dependent Ca^2+^ influx occurred (**Figure 6D**), that pharmacological induction of Ca^2+^ influx using calcimycin enabled vacuolar release (**Figure 6A**), and that blocking Ca^2+^ influx with nifedipine inhibited vacuolar release (**Figure 6G**). Calcium can function as a potent second messenger involved in numerous cellular processes (*49–51*). Changes in intracellular Ca^2+^ levels have been shown during infection with other pathogens able to survive inside cells, including *L. monocytogenes* (*52*). Moreover, Ca^2+^ has been implicated in cellular egress of *Chlamydia* species, potentially by influencing the activity of cellular proteases (*53–55*). Possibly also relevant, Ca^2+^ is involved in regulating autophagy, a process that can be blocked by *P. aeruginosa* (*49*, *56–58*). Manipulation of intracellular Ca^2+^ by the T3SS machinery might represent a previously unknown mechanism of blocking autophagy and therefore also bacterial clearance.

Results with the calcium channel blocker nifedipine showed two mechanisms by which the *P. aeruginosa* T3SS enables Ca^2+^ influx, one involving the T3SS exotoxins dependent on cellular Ca^2+^-channels and another involving the T3SS needle/translocon. In the absence of the T3SS exotoxins the remaining Ca^2+^ influx was independent of cellular Ca^2+^-channels, not surprising given that the T3SS translocon can form pores in host cell membranes (*31*). Thus, we were unable use a calcium channel blocker to explore the relationship between Ca^2+^ influx and vacuole release for the T3SS translocon proteins. However, we did find a cause-and-effect relationship between Ca^2+^ influx and vacuole release for the T3SS exotoxins. Moreover, pharmacological induction of Ca^2+^ influx alone triggered vacuole release suggesting generality for Ca^2+^ influx as a mediator. Testing this more directly for T3SS translocon-mediated Ca^2+^ influx will necessitate developing a strategy to inhibit it in a future study.

At first glance, some outcomes might suggest alternative mechanisms. For example, mutants constitutively expressing the T3SS caused more cell death than either wild-type or T3SS knockout mutants, hinting at involvement of cell death in the mechanism. However cell death occurred around 4-6 hour after vacuolar release suggesting it followed rather than being involved (**Figure 3B/D**). In fact, cell death after vacuole escape is expected because the presence of pattern recognition receptors in the cytoplasm can trigger inflammasome activation and subsequent cell death (*59*, *60*), which we showed specifically for *P. aeruginosa* inside corneal epithelial cells (*61*), a response the bacteria can modify using T3SS exotoxin ExoS (*61*). Also expected, were the similar phenotypes demonstrated by the Δ*pscC* (T3SS needle) and Δ*popBD* (T3SS translocon) mutants because needle mutants cannot assemble the translocon (*62*). Also worth discussing, is why the Δ*exsE* T3SS^ON^ mutant constitutively expressing the T3SS invaded as efficiently as wild-type and T3SS^OFF^ mutants (**Figure 3A**) whilst that the T3SS has potential for antiphagocytic activity (*23*, *38*). Possibilities include functional delays in correctly directed needle assembly, translocon insertion, and/or exotoxin secretion, and/or for them to impact the biology of the host cell in a manner that prevents phagocytosis. Understanding the timing of T3SS involvement in modulating internalization warrants further investigation.

For the *in vivo* experiments we infected live animals, used novel imaging methods that specifically distinguish intracellular bacteria, imaged the eye intravitally (i.e. in unfixed vital mouse eyes) to reduce the possibility of artifacts, and were able to observe the infection process with subcellular resolution. These methods enabled the study of cellular microbiology *in vivo*, including the potential for studying individual bacteria during infection temporally in addition to spatially. Using these methods, we showed that T3SS^OFF^ mutants (Δ*exsA*) used alone consistently localized in circular regions within cells in small foci. When T3SS^ON^ mutants (Δ*exsE*) were also present, the T3SS^OFF^ intracellular bacteria instead localized to more diffused punctate regions in the cell and were found across a larger area within the tissue (**Figure 7A/B**, **Supp Video 1**). While we remain unable to detect membranes around vacuoles within live eyes (only the plasma membrane), prior studies have used transmission electron microscopy to show that bacteria can reside in vacuoles *in vivo* (*63*). Thus, dispersal of vacuole-like structures in favor of smaller particles when T3SS^ON^ bacteria were used for co-infection likely represented cytoplasmic spreading populations, as found in *in vitro* co-infections. Further studies will be needed to explore if this phenotype switch seen for the T3SS^OFF^ mutants *in vivo* in the presence of mutants constitutively expressing the T3SS depends on the T3SS translocon pore as it does with cultured cells and if there is an impact on disease severity and outcome.

A caveat of the *in vivo* experiments was that a 1 hour infection was insufficient to enable bacteria to become intracellular as occurred *in vitro* in cell culture experiments. This was likely due to the anatomy and physiology *in vivo* creating a lack of synchrony that cannot be controlled for. Thus, eyes were infected for 16 hours before antibiotic was added for the final 4 hours. For that reason, some bacteria might have exited one cell and infected another before the extracellular bacteria were killed. Caveats such as this are routinely associated with *in vivo* experimentation and necessitate tandem *in vitro* experimentation as was done here, which also enabled inclusion of human cells. Notwithstanding, this caveat would have impacted all groups of mice and therefore cannot explain the differences found in the location of intracellular bacteria.

The significance of these findings toward patient management includes potential relevance to unpredictable or ineffective responses to antibiotics despite antibiotic sensitivity *in vitro*. Some antibiotics commonly used in a clinical setting are not cell-permeable and kill only extracellular bacteria, examples including the aminoglycosides. These antibiotics could therefore retain bacteria inside vacuoles that would otherwise be released by extracellular bacteria. The significance to disease pathogenesis is likely to depend on the infection site and other *in vivo* factors, e.g. regularly sloughing epithelial cell multilayers versus permanent epithelial cell monolayers, or superficial tissues versus enclosed organs. Importantly, our previously published work showed that vacuolar *P. aeruginosa* additionally resist cell-permeable antibiotics such as the fluoroquinolones, also commonly used for *P. aeruginosa* infections (*16*).

Here we showed that a simple change in an *in vitro* assay, shortening the time available for infection before killing extracellular bacteria, can drastically change the phenotype observed inside a cell. *In vivo* conditions surrounding infection can vary among patients, in the same patient within the infected tissue, and across the infection time span - including the timing and approach to treatment. This highlights the importance of assay design and studying phenotypes *in vivo*, while also challenging the classification of bacteria as either intracellular or extracellular.

In summary, the results of this study add yet another layer of complexity to our understanding of bacterial pathogenesis by showing that after diversifying into multiple subpopulations varying in location and gene expression, bacteria can collaborate across cell membrane barriers to promote pathogenesis. Release of vacuolar *P. aeruginosa* by extracellular *P. aeruginosa* requires cooperating across two membrane barriers. In this way, *P. aeruginosa* provides another interesting example of the value of diversity and collaboration with potential relevance to other bacterial pathogens and beyond.

## Supporting information

Schator et al Video 1

## Acknowledgements

Thanks to Dr. Arne Rietsch (Case Western Reserve University, OH, USA) for his valuable input and providing the pEXG2-Δ*exsE* plasmid, and to Dr. Timothy Yahr (University of Iowa, IA, USA) who provided the pJNE05 plasmid. Thanks also to Dr. Alain Filloux (Imperial College London, UK) for originally providing *P. aeruginosa* wild-type PAO1F, and to Dr. Danielle Robertson (University of Texas South Western, TX, USA) for originally providing telomerase-immortalized human corneal epithelial cells.

## Author contributions

DS, NGK, DE, and SF designed the experiments; DS, NGK, SJUC, and TKJ performed the experiments; DS, NGK, DE, and SF analyzed and interpreted the data; EJ and BES wrote the vacuole analysis macro; DS, DE and SF wrote the manuscript; DE and SF supervised the study.

## Methods

### Bacterial and cell culture

*Pseudomonas aeruginosa* strain PAO1F was used throughout the study and a list of mutants and plasmids used can be found in Table 1. Bacterial cultures were grown in Trypticase Soy broth (TSB) (Hardy Diagnostics, Santa Maria, CA, USA) or on Trypticase Soy agar (TSA) (Hardy Diagnostics) supplemented with Gentamicin (100 µg/ml) (Thermo Fisher Scientific, Hayward, CA, USA). Corneal epithelial cells (hTCEpi) (*64*) were maintained in KGM2 media (Lonza) in a 5% CO_2_ incubator (humidified) at 37 °C, supplemented with 1 mM of CaCl_2_ to induce differentiation before infection.

**Table 1:**
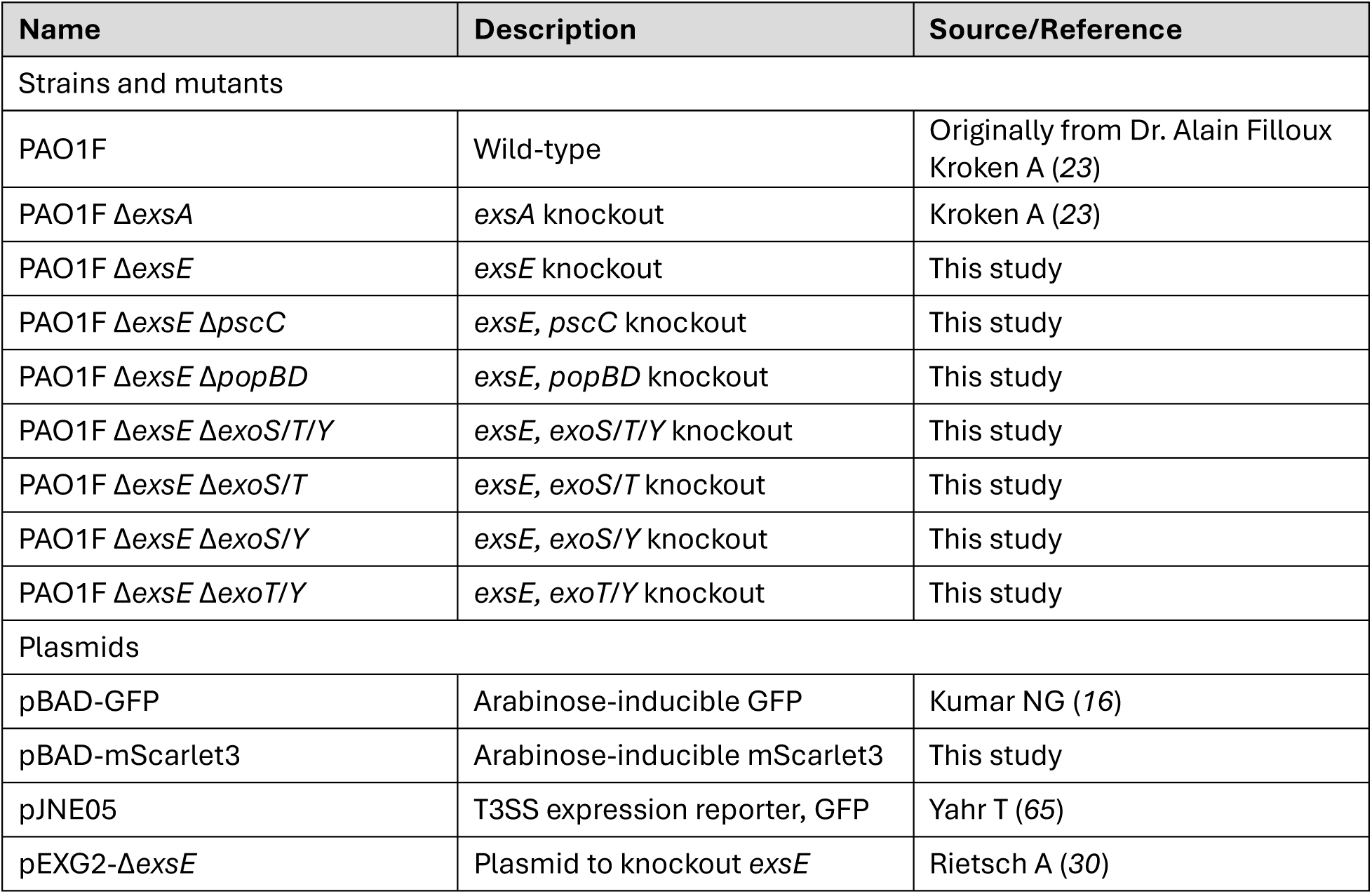
*P. aeruginosa* strains, mutants, and plasmids used in this study.

| Name | Description | Source/Reference |
| --- | --- | --- |
| Strains and mutants |  |  |
| PAO1F | Wild-type | Originally from Dr. Alain Filloux Kroken A (23) |
| PAO1F ΔexsA | exsA knockout | Kroken A (23) |
| PAO1F ΔexsE | exsE knockout | This study |
| PAO1F ΔexsE ΔpscC | exsE, pscC knockout | This study |
| PAO1F ΔexsE ΔpopBD | exsE, popBD knockout | This study |
| PAO1F ΔexsE ΔexoS/T/Y | exsE, exoS/T/Y knockout | This study |
| PAO1F ΔexsE ΔexoS/T | exsE, exoS/T knockout | This study |
| PAO1F ΔexsE ΔexoS/Y | exsE, exoS/Y knockout | This study |
| PAO1F ΔexsE ΔexoT/Y | exsE, exoT/Y knockout | This study |
| Plasmids |  |  |
| pBAD-GFP | Arabinose-inducible GFP | Kumar NG (16) |
| pBAD-mScarlet3 | Arabinose-inducible mScarlet3 | This study |
| pJNE05 | T3SS expression reporter, GFP | Yahr T (65) |
| pEXG2-ΔexsE | Plasmid to knockout exsE | Rietsch A (30) |

### Mutant construction

Isogenic mutants (Table 1) were constructed through homologues recombination using the pEXG2 plasmid backbone, containing the 500 bp regions up- and downstream of the gene to be deleted and bacterial conjugation using *Escherichia coli* SM10 (*66*). In short, *E. coli* SM10 containing the knockout plasmid and *P. aeruginosa* were grown separately in 3 ml TSB liquid culture overnight. The next morning, 3 ml of fresh TSB were added to *P. aeruginosa*, and bacteria were incubated at 42 °C for approximately 4 hours. The SM10 culture was used to inoculate fresh media in a 1:50 dilution and the bacteria were incubated at 37 °C, shaking for 4 hours. After the SM10 culture reached the exponential growth phase (OD_600_ of 0.2-0.4), they were mixed with *P. aeruginosa* at a ratio of 3:1 (*E. coli*:*Pa*), centrifuged at 10,000 x g for 4 min, then resuspended in 50 µl of Luria broth (LB), spotted on LB plates and incubated at 37 °C overnight. The next day, the spot of bacteria is scrapped off, resuspended in sterile PBS, then plated onto Vogel-Bonner minimal media (VBMM) agar supplemented with gentamicin (100 µg/ml). Single colonies were picked and used to inoculate 1 ml LB cultures for 4 hours at 37 °C. Those cultures are then plated for counterselection on Yeast-extract/Tryptone (YT) plates, containing 15 % sucrose. Single colonies from the counterselection plates were patched on TSA and TSA with Gentamicin (100 µg/ml) plates to check for plasmid excision (i.e., no growth on TSA with gentamicin). Mutations were then verified by colony PCR and fragment sequencing.

### Infection assays and timelapse imaging

Corneal cells (hTCEpi) were seeded in a 24-well glass bottom dish (MatTek) and 1 mM CaCl_2_ was added to the cell culture media to induce cell differentiation. Cells were incubated at 37 °C overnight (humidified) in a 5 % CO_2_ incubator. Bacteria were grown on TSA plates, containing gentamicin (100 µg/ml) for plasmid retention. Bacteria were collected from the plate using a sterile loop and were resuspended in PBS. OD_600_ was used to calculate bacterial numbers for infections. Before infection, Hoechst 33342 (ImmunoChemistry Technologies, Davis, CA, USA) was added to visualize cell nuclei. At 1 or 3 hours post-infection the cell culture media was replaced with fresh media containing amikacin (200 µg/ml) and CaCl_2_ (1 mM). After another 30 minutes, the media was replaced again with cell culture media containing amikacin (200 µg/ml), CaCl_2_ (1 mM) and 0.5 % L-arabinose (to induce plasmid-encoded fluorophore expression) as well as either propidium iodide (0.75 µg/ml) (PI) (ImmunoChemistry Technologies) or DRAQ7 (2 µl/ml) (Thermo Fisher Scientific) to visualize cell permeabilization. Depending on the experiment, different compounds were also added at this step (e.g., saponin [Thermo Fisher Scientific, 0.03 mg/ml], valinomycin [Sigma Aldrich {Burlington, MA, USA}, 500 nM], calcimycin [Sigma Aldrich, 1 µM]). The glass bottom dish was set up in a wide-field microscope for timelapse imaging. Images were acquired using a Nikon Ti-E inverted wide-field fluorescence microscope, paired with a Lumencor SpectraX illumination source. To observe live samples, cells were incubated in an Okolab Uno-combined controller stage top incubation chamber to ensure consistent heat, humidity, and 5% CO_2_. For timelapse imaging a CFI Plan Apo Lambda 40X NA 0.95 air objective, equipped with differential interference contrast (DIC), was used. Nikon Perfect Focus hardware maintained focal planes throughout imaging. For each condition eight fields of view were selected for timelapse studies prioritizing areas devoid of debris using DIC. GFP/mScarlet3 observation was deferred until completion to prevent bias.

### Growth curves and reporter assays

In a 96-well plate, 100 µl of TSB was added per well. For the reporter assays, the media was supplemented with 1% glycerol, 100 µM mono sodium glutamate (MSG) and 2 mM EGTA (EGTA was not added for negative controls). Approximately 10^6^ cfu bacteria were added to each well and growth (OD_600_) and fluorescence of the T3SS reporter (pJNE05) were tracked using a BioTek Synergy HTX multimode reader at 37°C with measurements taken every 30 minutes. The plate was shaken during the measurements and the growth was followed for 20 hours.

### T3SS activity spotting assay

*P. aeruginosa* containing a plasmid for detection of T3SS expression (pJNE05) were used to inoculate 2 ml TSB cultures containing 100 µg/ml gentamicin which were incubated overnight at 37 °C. The next day, these cultures were used at a dilution of 1:10,000 to inoculate induction cultures (see Growth curves and reporter assays). After 20 hours of growth in a shaking incubator at 37 °C, 100 µl of the bacterial suspension was mixed with 100 µl of 4% paraformaldehyde and bacteria fixed for 15 min at room temperature. Samples were centrifuged at 10,000 x g for 10 min and bacterial pellets were resuspended in 1 ml PBS containing 3 µl/ml of propidium iodide (to stain bacteria). After 1 hour of incubation at room temperature, the samples were centrifuged again, and pellets resuspended in 1 ml of purified water. 100 µl of this suspension was spotted on glass cover slips and dried at 37 °C for 2 hours. Cover slips were then mounted on glass slides using ProLong Diamond Antifade mountant (Thermo Fisher Scientific). Samples were cured over night at room temperature and then imaged using a Nikon Ti-E inverted wide-field fluorescence microscope as above, but at 60x magnification (oil immersion, NA 1.4). GFP-spot intensity was measured with a macro in ImageJ. The full macro can be found online (DOI: 10.5281/zenodo.14797030).

### Time lapse image analysis

For image analysis ImageJ (version 2.14.0/1.54f) was used. A macro was developed to measure vacuole numbers, infection rates and cell death. In short, the macro contains several critical steps to analyze bacteria containing vacuoles. First, the macro marks and counts all the nuclei, based on their Hoechst signals and cross-checks these results for cell death staining, excluding all nuclei that show a cell death signal. It then takes those spots and enlarges them to a certain degree and checks their area for signal of the bacterial fluorophore (since we previously showed that the vacuoles are perinuclear (*13*)). The next step entails several measurements of the bacterial signal to determine if it fulfills the criteria for vacuoles (the signal needs to have a minimum area and a certain degree of circularity). The macro counts all of these events in every field of view and generates an Excel sheet with the data output. The full macro can be found online (DOI: 10.5281/zenodo.14797030).

### Calcium sensor experiments

Corneal cells (hTCEpi) were seeded onto a 24-well glass bottom dish (MatTek) with 1 mM CaCl_2_ added as above. For experiments using the calcium-channel blocker nifedipine (Thermo Fisher Scientific), 10 µM of nifedipine was added to the cells at the same time. Bacteria with the pBAD-mScarlet3 plasmid were grown on TSA plates, containing gentamicin. Bacteria are collected from the plate using a sterile loop and are resuspended in PBS. OD_600_ is used to calculate bacterial numbers for infections. Before infection, Hoechst 33342 (ImmunoChemistry Technologies) was added to visualize cell nuclei. At 1 hour post-infection the cell culture media was replaced with fresh media containing amikacin (200 µg/ml) and CaCl_2_ (1 mM). After another 30 minutes, the media was replaced again with cell culture media with amikacin (200 µg/ml), CaCl_2_ (1 mM) and 0.5 % L-arabinose (to induce plasmid-encoded fluorophore expression) as well as DRAQ7 (0.6 µM) (Thermo Fisher Scientific) to visualize cell permeabilization. At four hours post-infection, the cells were washed once with PBS then incubated with cell culture media with Fluo-5F AM (Thermo Fisher Scientific) at 2 µM for 30 minutes at 37 °C. Then, cells were washed again with PBS and fresh KGM2 was added to the cells. After a further incubation for 30 minutes at 37 °C, images are taken using the widefield microscope at 40x magnification. Images were analyzed using an ImageJ macro which was made public online (DOI: 10.5281/zenodo.14797030).

### Probability calculations

Probability calculation was used to predict the rate of co-infection of wild-type or ΔexsA with Δ*exsE* in the same cell. The formula for both events occurring at the same time was as follows:

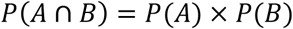

Here, P(A) is the infection rate of wild-type bacteria or the Δ*exsA* strain and P(B) is the infection rate of the Δ*exsE* mutant. The calculated (predicted) values are then compared to the observed values from experiments. Next, the predicted rate of co-infection with Δ*exsE* mutant bacteria within the wild-type or Δ*exsA* infected cells was calculated. To obtain the predicted rate of co-infection within the wild-type or Δ*exsA* infected cells, a conditional probability calculation was performed using the following formula:

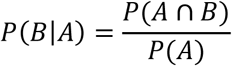

In our case, P(A∩B) is the observed rate of co-infection of either wild-type or Δ*exsA* with the Δ*exsE* mutant in the same cell in the overall population and P(A) is the infection rate of either wild-type or Δ*exsA* alone. The resulting value is the predicted probability of co-infection with the Δ*exsE* mutant within the wild-type or Δ*exsA* infected cells. Our hypothesis was that co-infection of wild-type or Δ*exsA* infected cells with the Δ*exsE* mutant was necessary for their vacuolar release. If this was the case, the predicted value of co-infection within these populations should be the same as the observed value of vacuolar release in the same populations. These calculations were performed for different biological replicates and then plotted with standard deviation on grouped bar graphs comparing the calculated (predicted) values to the values observed in the experiments.

### Murine corneal scratch infection model

All procedures involving animals were carried out in accordance with the standards established by the Association for the Research in Vision and Ophthalmology, under protocol AUP-2019-06-12322-1, approved by the Animal Care and Use Committee, University of California Berkeley. This protocol adheres to Public Health Service policy on the humane care and use of laboratory animals and the guide for the care and use of laboratory animals. For the *in vivo* imaging 8- to 16-week old mice (male and female) ROSA^mT/mG^ mice were used for their expression of dtTomato on their cell membranes. The mice were anesthetized by ketamine-dexmedetomidine injection (ketamine, 80-100 mg/kg; dexmedetomidine, 0.25 to 0.5 mg/kg). For one eye, the cornea was scratched three times in parallel lines using a 26 G needle. The scratched eyes were inoculated with 5 µl of bacterial suspension (either ∼10^10^ [90%+10%] or ∼10^9^ [10%] CFU/ml). Bacteria with a plasmid for inducible GFP-expression (pBAD-GFP) and bacteria without the plasmid were mixed in different ratios: 10% Δ*exsE*-GFP (50 µl Δ*exsE* [GFP expressing] + 450 µl PBS), 90% Δ*exsA*-GFP+10% Δ*exsE* (450 µl Δ*exsA* [GFP expressing] + 50 µl Δ*exsE* [no GFP]), 90% Δ*exsA*-GFP+10% Δ*exsA* (450 µl Δ*exsA* [GFP expressing] + 50 µl Δ*exsA* [no GFP]). After scratch and inoculation, the mice were injected with anesthesia-reversal (atipamezole hydrochloride, 2.5-5 mg/kg). The next day (16 hours post-infection), mice were again anesthetized using ketamine-dexmedetomidine injection. Infected eyes were then treated every hour with RPMI media + amikacin (300 µg/ml) for two hours to kill all extracellular bacteria, followed by treatment with RPMI media + amikacin (300 µg/ml) + L-arabinose (5%), for two more hours to induce bacterial GFP-expression. After treatment, the mice were sacrificed by ketamine-xylazine injection (ketamine, 80-100 mg/kg; xylazine, 5-10 mg/kg) followed by cervical dislocation. Eyes were enucleated then imaged live without fixation by confocal microscopy (Olympus FluoView) using a 60x water immersion objective, with an argon laser, a pinhole size of 105 µm, and lasers set for FITC (excitation 488 nm, emission 500-545 nm) and TRITC (excitation 559 nm, emission 570-670 nm).

### Corneal scratch image analysis

To determine GFP distribution, images were analyzed using ImageJ (version 2.14.0/1.54f). In brief, a 100×100 µm area was chosen based on location of infection foci and background subtraction was performed. A GFP histogram was generated for the measured area based on the maximum intensity projection and the values normalized to the maximum signal within the analyzed region. These steps were repeated for each image and the data was accumulated and plotted on a histogram, including LOWESS curves.

To determine GFP aggregate sizes and the percent infected cells, Imaris software (v 10.1) was used. A batch analysis macro was created to standardize image analysis across experiments. In brief, background subtraction (GFP) and baseline subtraction (dtTomato) was performed, followed by the cell + vesicle function (bacterial aggregates were considered vesicles for the analysis). Based on the dtTomato signal of cell membranes, the software automatically determined the localization and outline of each cell. Bacterial signal within the cells was differentiated into two types, “single bacteria” – GFP signal with a diameter of around 1.5 µm – and “aggregates”, particles with a diameter of 5 µm and above. The macro then measures the volume of every detected particle and determines in which cell it was located. This data was then used to determine the percent infected cells as well as the average size of bacterial aggregates in the different infection conditions. The batch analysis settings have been made publicly available online (DOI: 10.5281/zenodo.14797030).

### Statistics

Statistical analyses were performed using GraphPad Prism 10. Data were shown as mean ± SEM (for analyses based on images, with them representing averages for several independent experiments, each containing 8 fields of view) or mean ± SD for 3-8 independent experiments. For column analyses with one categorical variable, a Student’s t-test was performed. For the grouped analysis of two or more groups with one categorical variable, a One-way ANOVA with multiple comparisons (Tukey’s post hoc analysis) was performed. For the grouped analysis of two or more groups with two categorical variables, a Two-way ANOVA with multiple comparisons (Tukey’s post hoc analysis) was performed. For every analysis P < 0.05=*, P < 0.01=**, P < 0.005=***, and P < 0.001=****. P values less than 0.05 were considered significant.

---

**Video 1: Representative *in vivo* confocal microscopy videos of infected murine corneal epithelium**

Representative *in vivo* confocal microscopy videos of murine corneal epithelium (ROSAmT/mG mice, dtTomato cell membranes, red) from apical to basal side of the corneal epithelium infected with different combinations of fluorescent (green) and non-fluorescent bacteria (Δ*exsA*-GFP+10% Δ*exsA*, 90% Δ*exsA*-GFP+10% Δ*exsE*, 10% Δ*exsE*-GFP). Images were taken 20 hours post-infection using the murine scratch-infection model at infection foci within the corneal epithelium using a 60x water-immersion objective. The videos first show a 3-D view of the infected corneal epithelium (apical and basal side are indicated in the video), followed by an orthogonal projection of the stack moving from the apical to the basal side and back.

## References

1. A. C. Blanchard, V. J. Waters, Opportunistic Pathogens in Cystic Fibrosis: Epidemiology and Pathogenesis of Lung Infection. J Pediatric Infect Dis Soc 11, S3–S12 (2022).

2. L. K. Branski, A. Al-Mousawi, H. Rivero, M. G. Jeschke, A. P. Sanford, D. N. Herndon, Emerging infections in burns. Surg Infect (Larchmt) 10, 389–397 (2009).

3. S. M. J. Fleiszig, A. R. Kroken, V. Nieto, M. R. Grosser, S. J. Wan, M. M. E. Metruccio, D. J. Evans, Contact lens-related corneal infection: Intrinsic resistance and its compromise. Prog Retin Eye Res 76, 100804 (2020).

4. F. Stapleton, E. Pearlman, C. Tam, A. Campolo, R. Pifer, P. Shannon, M. Crary, Microbial Adherence to Contact Lenses and Pseudomonas aeruginosa as a Model Organism for Microbial Keratitis. Pathogens 2022, Vol. 11, Page 1383 11, 1383 (2022).

5. D. C. Wu, W. W. Chan, A. I. Metelitsa, L. Fiorillo, A. N. Lin, Pseudomonas Skin Infection. American Journal of Clinical Dermatology 2011 12:3 12, 157–169 (2012).

6. C. C. Szeto, P. K. T. Li, Peritoneal dialysis–associated peritonitis. Clinical Journal of the American Society of Nephrology 14, 1100–1105 (2019).

7. S. J. T. Wardell, J. Gauthier, L. W. Martin, M. Potvin, B. Brockway, R. C. Levesque, I. L. Lamont, Genome evolution drives transcriptomic and phenotypic adaptation in Pseudomonas aeruginosa during 20 years of infection. Microb Genom 7 (2021).

8. E. Rossi, R. La Rosa, J. A. Bartell, R. L. Marvig, J. A. J. Haagensen, L. M. Sommer, S. Molin, H. K. Johansen, Pseudomonas aeruginosa adaptation and evolution in patients with cystic fibrosis. Nat Rev Microbiol 19, 331–342 (2021).

9. Y. Hilliam, M. P. Moore, I. L. Lamont, D. Bilton, C. S. Haworth, J. Foweraker, M. J. Walshaw, D. Williams, J. L. Fothergill, A. De Soyza, C. Winstanley, Pseudomonas aeruginosa adaptation and diversification in the non-cystic fibrosis bronchiectasis lung. European Respiratory Journal 49 (2017).

10. D. Ruhluel, L. Fisher, T. E. Barton, H. Leighton, S. Kumar, P. Amores Morillo, S. O’Brien, J. L. Fothergill, D. R. Neill, Secondary messenger signalling influences Pseudomonas aeruginosa adaptation to sinus and lung environments. ISME J 18, 65 (2024).

11. A. Weimann, A. M. Dinan, C. Ruis, A. Bernut, S. Pont, K. Brown, J. Ryan, L. Santos, L. Ellison, E. Ukor, A. P. Pandurangan, S. Krokowski, T. L. Blundell, M. Welch, B. Blane, K. Judge, R. Bousfield, N. Brown, J. M. Bryant, I. Kukavica-Ibrulj, G. Rampioni, L. Leoni, P. T. Harrison, S. J. Peacock, N. R. Thomson, J. Gauthier, J. L. Fothergill, R. C. Levesque, J. Parkhill, R. A. Floto, Evolution and host-specific adaptation of Pseudomonas aeruginosa. Science (1979) 385 (2024).

12. A. Weimann, A. M. Dinan, C. Ruis, A. Bernut, S. Pont, K. Brown, J. Ryan, L. Santos, L. Ellison, E. Ukor, A. P. Pandurangan, S. Krokowski, T. L. Blundell, M. Welch, B. Blane, K. Judge, R. Bousfield, N. Brown, J. M. Bryant, I. Kukavica-Ibrulj, G. Rampioni, L. Leoni, P. T. Harrison, S. J. Peacock, N. R. Thomson, J. Gauthier, J. L. Fothergill, R. C. Levesque, J. Parkhill, R. A. Floto, Evolution and host-specific adaptation of Pseudomonas aeruginosa. Science 385, eadi0908 (2024).

13. A. A. Angus, A. A. Lee, D. K. Augustin, E. J. Lee, D. J. Evans, S. M. J. Fleiszig, Pseudomonas aeruginosa induces membrane blebs in epithelial cells, which are utilized as a niche for intracellular replication and motility. Infect Immun 76, 1992–2001 (2008).

14. A. L. Jolly, D. Takawira, O. O. Oke, S. A. Whiteside, S. W. Chang, E. R. Wen, K. Quach, D. J. Evans, S. M. J. Fleiszig, Pseudomonas aeruginosa-Induced Bleb-Niche Formation in Epithelial Cells Is Independent of Actinomyosin Contraction and Enhanced by Loss of Cystic Fibrosis Transmembrane-Conductance Regulator Osmoregulatory Function. mBio 6 (2015).

15. A. R. Kroken, N. G. Kumar, T. L. Yahr, B. E. Smith, V. Nieto, H. Horneman, D. J. Evans, S. M. J. Fleiszig, Exotoxin S secreted by internalized Pseudomonas aeruginosa delays lytic host cell death. PLoS Pathog 18 (2022).

16. N. G. Kumar, V. Nieto, A. R. Kroken, E. Jedel, M. R. Grosser, M. E. Hallsten, M. M. E. Mettrucio, T. L. Yahr, D. J. Evans, S. M. J. Fleiszig, Pseudomonas aeruginosa Can Diversify after Host Cell Invasion to Establish Multiple Intracellular Niches. mBio 13 (2022).

17. S. M. J. Fleiszig, T. S. Zaidi, E. L. Fletcher, M. J. Preston, G. B. Pier, Pseudomonas aeruginosa invades corneal epithelial cells during experimental infection. Infect Immun 62, 3485–3493 (1994).

18. S. M. J. Fleiszig, T. S. Zaidi, M. J. Preston, M. Grout, D. J. Evans, G. B. Pier, Relationship between cytotoxicity and corneal epithelial cell invasion by clinical isolates of Pseudomonas aeruginosa. Infect Immun 64, 2288–2294 (1996).

19. S. M. J. Fleiszig, T. S. Zaidi, G. B. Pier, Pseudomonas aeruginosa invasion of and multiplication within corneal epithelial cells in vitro. Infect Immun 63, 4072–4077 (1995).

20. A. Leoni Swart, B. J. Laventie, R. Sütterlin, T. Junne, L. Lauer, P. Manfredi, S. Jakonia, X. Yu, E. Karagkiozi, R. Okujava, U. Jenal, Pseudomonas aeruginosa breaches respiratory epithelia through goblet cell invasion in a microtissue model. Nature Microbiology 2024 9:7 9, 1725–1737 (2024).

21. K. Malet, E. Faure, D. Adam, J. Donner, L. Liu, S. J. Pilon, R. Fraser, P. Jorth, D. K. Newman, E. Brochiero, S. Rousseau, D. Nguyen, Intracellular Pseudomonas aeruginosa within the Airway Epithelium of Cystic Fibrosis Lung Tissues. Am J Respir Crit Care Med 209, 1453–1462 (2024).

22. P. Garai, L. Berry, M. Moussouni, S. Bleves, A. B. Blanc-Potard, Killing from the inside: Intracellular role of T3SS in the fate of pseudomonas aeruginosa within macrophages revealed by mgtC and oprF mutants. PLoS Pathog 15 (2019).

23. A. R. Kroken, C. K. Chen, D. J. Evans, T. L. Yahr, S. M. J. Fleiszig, The impact of ExoS on Pseudomonas aeruginosa internalization by epithelial cells is independent of fleQ and correlates with bistability of type three secretion system gene expression. mBio 9 (2018).

24. V. Hritonenko, D. J. Evans, S. M. J. Fleiszig, Translocon-Independent Intracellular Replication by Pseudomonas aeruginosa Requires the ADP-Ribosylation Domain of ExoS. Microbes and infection / Institut Pasteur 14, 1366 (2012).

25. S. R. Heimer, D. J. Evans, M. E. Stern, J. T. Barbieri, T. Yahr, S. M. J. Fleiszig, Pseudomonas aeruginosa Utilizes the Type III Secreted Toxin ExoS to Avoid Acidified Compartments within Epithelial Cells. PLoS One 8, e73111 (2013).

26. A. A. Angus, D. J. Evans, J. T. Barbieri, S. M. J. Fleiszig, The ADP-ribosylation domain of Pseudomonas aeruginosa ExoS is required for membrane bleb niche formation and bacterial survival within epithelial cells. Infect Immun 78, 4500–4510 (2010).

27. J. Fredlund, J. C. Santos, V. Stévenin, A. Weiner, P. Latour-Lambert, K. Rechav, A. Mallet, J. Krijnse-Locker, M. Elbaum, J. Enninga, The entry of Salmonella in a distinct tight compartment revealed at high temporal and ultrastructural resolution. Cell Microbiol 20, e12816 (2018).

28. N. Dasgupta, A. Ashare, G. W. Hunninghake, T. L. Yahr, Transcriptional Induction of the Pseudomonas aeruginosa Type III Secretion System by Low Ca2+ and Host Cell Contact Proceeds through Two Distinct Signaling Pathways. Infect Immun 74, 3334 (2006).

29. E. Faure, J. B. Mear, K. Faure, S. Normand, A. Couturier-Maillard, T. Grandjean, V. Balloy, B. Ryffel, R. Dessein, M. Chignard, C. Uyttenhove, B. Guery, P. Gosset, M. Chamaillard, E. Kipnis, Pseudomonas aeruginosa type-3 secretion system dampens host defense by exploiting the NLRC4-coupled inflammasome. Am J Respir Crit Care Med 189, 799–811 (2014).

30. A. Rietsch, I. Vallet-Gely, S. L. Dove, J. J. Mekalanos, ExsE, a secreted regulator of type III secretion genes in Pseudomonas aeruginosa. Proc Natl Acad Sci U S A 102, 8006–8011 (2005).

31. L. Dortet, C. Lombardi, F. Cretin, A. Dessen, A. Filloux, Pore-forming activity of the Pseudomonas aeruginosa type III secretion system translocon alters the host epigenome. Nature Microbiology 2018 3:3 3, 378–386 (2018).

32. M. A. Hamon, P. Cossart, K+ efflux is required for histone H3 dephosphorylation by Listeria monocytogenes listeriolysin o and otherpore-forming toxins. Infect Immun 79, 2839–2846 (2011).

33. S. Zhang, Y. Zheng, S. Chen, S. Huang, K. Liu, Q. Lv, Y. Jiang, Y. Yuan, Suilysin-induced Platelet-Neutrophil Complexes Formation is Triggered by Pore Formation-dependent Calcium Influx. Scientific Reports 2016 6:1 6, 1–11 (2016).

34. R. Fiser, J. Masin, L. Bumba, E. Pospisilova, C. Fayolle, M. Basler, L. Sadilkova, I. Adkins, J. Kamanova, J. Cerny, I. Konopasek, R. Osicka, C. Leclerc, P. Sebo, Calcium Influx Rescues Adenylate Cyclase-Hemolysin from Rapid Cell Membrane Removal and Enables Phagocyte Permeabilization by Toxin Pores. PLoS Pathog 8, e1002580 (2012).

35. J. Li, M. M. E. Metruccio, D. J. Evans, S. M. J. Fleiszig, Mucosal fluid glycoprotein DMBT1 suppresses twitching motility and virulence of the opportunistic pathogen Pseudomonas aeruginosa. PLoS Pathog 13, e1006392 (2017).

36. M. Ni, D. J. Evans, S. Hawgood, E. M. Anders, R. A. Sack, S. M. J. Fleiszig, Surfactant protein D is present in human tear fluid and the cornea and inhibits epithelial cell invasion by Pseudomonas aeruginosa. Infect Immun 73, 2147–2156 (2005).

37. N. A. McNamara, R. Andika, M. Kwong, R. A. Sack, S. M. J. Fleiszig, Interaction of Pseudomonas aeruginosa with Human Tear Fluid Components. Curr Eye Res 30, 517–525 (2005).

38. B. A. Cowell, D. Y. Chen, D. W. Frank, A. J. Vallis, S. M. J. Fleiszig, ExoT of cytotoxic Pseudomonas aeruginosa prevents uptake by corneal epithelial cells. Infect Immun 68, 403–406 (2000).

39. I. Alarcon, D. J. Evans, S. M. J. Fleiszig, The Role of Twitching Motility in Pseudomonas aeruginosa Exit from and Translocation of Corneal Epithelial Cells. Invest Ophthalmol Vis Sci 50, 2237 (2009).

40. M. Galle, S. Jin, P. Bogaert, M. Haegman, P. Vandenabeele, R. Beyaert, The Pseudomonas aeruginosa Type III Secretion System Has an Exotoxin S/T/Y Independent Pathogenic Role during Acute Lung Infection. PLoS One 7, e41547 (2012).

41. K. Nomura, K. Obata, T. Keira, R. Miyata, S. Hirakawa, K. ichi Takano, T. Kohno, N. Sawada, T. Himi, T. Kojima, Pseudomonas aeruginosa elastase causes transient disruption of tight junctions and downregulation of PAR-2 in human nasal epithelial cells. Respir Res 15, 1–13 (2014).

42. H. Yu, X. He, W. Xie, J. Xiong, H. Sheng, S. Guo, C. Huang, D. Zhang, K. Zhang, Elastase LasB of pseudomonas aeruginosa promotes biofilm formation partly through rhamnolipid-mediated regulation. Can J Microbiol 60, 227–235 (2014).

43. C. Schwarzer, H. Fischer, T. E. Machen, Chemotaxis and Binding of Pseudomonas aeruginosa to Scratch-Wounded Human Cystic Fibrosis Airway Epithelial Cells. PLoS One 11, e0150109 (2016).

44. P. Tielen, H. Kuhn, F. Rosenau, K. E. Jaeger, H. C. Flemming, J. Wingender, Interaction between extracellular lipase LipA and the polysaccharide alginate of Pseudomonas aeruginosa. BMC Microbiol 13, 1–12 (2013).

45. F. Rosenau, S. Isenhardt, A. Gdynia, D. Tielker, E. Schmidt, P. Tielen, M. Schobert, D. Jahn, S. Wilhelm, K. E. Jaeger, Lipase LipC affects motility, biofilm formation and rhamnolipid production in Pseudomonas aeruginosa. FEMS Microbiol Lett 309, 25–34 (2010).

46. K. Murphy, A. J. Park, Y. Hao, D. Brewer, J. S. Lam, C. M. Khursigaraa, Influence of O polysaccharides on biofilm development and outer membrane vesicle biogenesis in Pseudomonas aeruginosa PAO1. J Bacteriol 196, 1306–1317 (2014).

47. V. Finck-Barbançon, J. Goranson, L. Zhu, T. Sawa, J. P. Wiener-Kronish, S. M. J. Fleiszig, C. Wu, L. Mende-Mueller, D. W. Frank, ExoU expression by Pseudomonas aeruginosa correlates with acute cytotoxicity and epithelial injury. Mol Microbiol 25, 547–557 (1997).

48. B. H. Iglewski, J. Sadoff, M. J. Bjorn, E. S. Maxwell, Pseudomonas aeruginosa exoenzyme S: an adenosine diphosphate ribosyltransferase distinct from toxin A. Proceedings of the National Academy of Sciences 75, 3211–3215 (1978).

49. P. Sukumaran, V. N. Da Conceicao, Y. Sun, N. Ahamad, L. R. Saraiva, S. Selvaraj, B. B. Singh, Calcium Signaling Regulates Autophagy and Apoptosis. Cells 2021, Vol. 10, Page 2125 10, 2125 (2021).

50. P. Pinton, C. Giorgi, R. Siviero, E. Zecchini, R. Rizzuto, Calcium and apoptosis: ER-mitochondria Ca2+ transfer in the control of apoptosis. Oncogene 2008 27:50 27, 6407–6418 (2008).

51. D. E. Clapham, Calcium Signaling. Cell 131, 1047–1058 (2007).

52. T. Li, L. Kong, X. Li, S. Wu, K. S. Attri, Y. Li, W. Gong, B. Zhao, L. Li, L. E. Herring, J. M. Asara, L. Xu, X. Luo, Y. L. Lei, Q. Ma, S. Seveau, J. S. Gunn, X. Cheng, P. K. Singh, D. R. Green, H. Wang, H. Wen, Listeria monocytogenes upregulates mitochondrial calcium signalling to inhibit LC3-associated phagocytosis as a survival strategy. Nature Microbiology 2021 6:3 6, 366–379 (2021).

53. K. Hybiske, R. S. Stephens, Mechanisms of host cell exit by the intracellular bacterium Chlamydia. Proc Natl Acad Sci U S A 104, 11430–11435 (2007).

54. J. Scholz, G. Holland, M. Laue, S. Banhart, D. Heuer, *Chlamydia* -containing spheres are a novel and predominant form of egress by the pathogen *Chlamydia psittaci*. mBio, doi: 10.1128/MBIO.01288-24 (2024).

55. T. Yamashima, Ca2+-dependent proteases in ischemic neuronal death: A conserved ‘calpain–cathepsin cascade’ from nematodes to primates. Cell Calcium 36, 285–293 (2004).

56. X. Jin, C. Zhang, S. Lin, T. Gao, H. Qian, L. Qu, J. Yao, X. Du, G. Feng, Pec 1 of Pseudomonas aeruginosa Inhibits Bacterial Clearance of Host by Blocking Autophagy in Macrophages. ACS Infect Dis 10, 2741–2754 (2024).

57. A. Chakraborty, A. Kabashi, S. Wilk, L. G. Rahme, Quorum-Sensing Signaling Molecule 2-Aminoacetophenone Mediates the Persistence of Pseudomonas aeruginosa in Macrophages by Interference with Autophagy through Epigenetic Regulation of Lipid Biosynthesis. mBio 14 (2023).

58. L. Rao, I. De La Rosa, Y. Xu, Y. Sha, A. Bhattacharya, M. J. Holtzman, B. E. Gilbert, N. T. Eissa, Pseudomonas aeruginosa survives in epithelia by ExoS-mediated inhibition of autophagy and mTOR. EMBO Rep 22 (2021).

59. T. Grandjean, A. Boucher, M. Thepaut, L. Monlezun, B. Guery, E. Faudry, E. Kipnis, R. Dessein, The human NAIP-NLRC4-inflammasome senses the Pseudomonas aeruginosa T3SS inner-rod protein. Int Immunol 29, 377–384 (2017).

60. T. Wangdi, L. A. Mijares, B. I. Kazmierczak, In vivo discrimination of type 3 secretion system-positive and -negative Pseudomonas aeruginosa via a caspase-1-dependent pathway. Infect Immun 78, 4744–4753 (2010).

61. A. R. Kroken, K. A. Klein, P. S. Mitchell, V. Nieto, E. J. Jedel, D. J. Evans, S. M. J. Fleiszig, Intracellular replication of Pseudomonas aeruginosa in epithelial cells requires suppression of the caspase-4 inflammasome. mSphere 8 (2023).

62. E. Kundracik, J. Trichka, J. D. Aponte, A. Roistacher, A. Rietsch, PopB-PcrV Interactions Are Essential for Pore Formation in the Pseudomonas aeruginosa Type III Secretion System Translocon. mBio 13 (2022).

63. S. M. J. Fleiszig, T. S. Zaidi, E. L. Fletcher, M. J. Preston, G. B. Pier, Pseudomonas aeruginosa invades corneal epithelial cells during experimental infection. Infect Immun 62, 3485–3493 (1994).

64. D. M. Robertson, L. Li, S. Fisher, V. P. Pearce, J. W. Shay, W. E. Wright, H. D. Cavanagh, J. V. Jester, Characterization of Growth and Differentiation in a Telomerase-Immortalized Human Corneal Epithelial Cell Line. Invest Ophthalmol Vis Sci 46, 470–478 (2005).

65. M. L. Urbanowski, E. D. Brutinel, T. L. Yahr, Translocation of ExsE into Chinese hamster ovary cells is required for transcriptional induction of the Pseudomonas aeruginosa type III secretion system. Infect Immun 75, 4432–4439 (2007).

66. L. R. Hmelo, B. R. Borlee, H. Almblad, M. E. Love, T. E. Randall, B. S. Tseng, C. Lin, Y. Irie, K. M. Storek, J. J. Yang, R. J. Siehnel, P. L. Howell, P. K. Singh, T. Tolker-Nielsen, M. R. Parsek, H. P. Schweizer, J. J. Harrison, Precision-engineering the Pseudomonas aeruginosa genome with two-step allelic exchange. Nat Protoc 10, 1820–1841 (2015).

